# Wakeful rest benefits memory when materials can be rehearsed

**DOI:** 10.1101/449835

**Authors:** Peter R. Millar, David A. Balota

## Abstract

Wakeful rest is a brief (e.g., 10 minutes), quiet period of minimal stimulation, which has been shown to facilitate memory performance, compared to a distractor task. Researchers have argued that this benefit is driven by automatic consolidation during the wakeful rest period. However, prior studies have not fully ruled out a controlled rehearsal mechanism, which might also occur during wakeful rest. In the current study, we attempted to replicate the wakeful rest effect under conditions that more strictly limit the possibility of rehearsal. Across six experiments, we manipulated parameters of a standard wakeful rest paradigm, including the type of target materials (word lists or abstract shapes), intentionality of encoding (incidental or intentional), and final retrieval delay (immediate or delayed). Additionally, we tested both younger and older adults to test whether these effects are consistent across the adult lifespan. Importantly, we observed the expected wakeful rest memory benefit in recall for verbal targets, which are easily rehearseable, but not for abstract shapes, which cannot be readily rehearsed. This pattern occurred in both younger and older adults. These results place constraints on the generalizability of wakeful rest memory benefits and suggest that the effect may be at least partly driven by rehearsal processes, rather than an automatic consolidation process.

---

Memory consolidation has been viewed as involving neurological processes, ranging from the molecular to the systems levels, acting to stabilize memory traces over time (for review, see 1-5; but also see 6,7). These consolidation processes are theorized to ultimately transfer the storage of a memory trace from hippocampal to neocortical areas, resulting in a stronger long-term trace that is less prone to forgetting. Evidence for consolidation, comes from a wide, interdisciplinary range of sources including studies of amnestic individuals (e.g., 9,10), animal lesion experiments (e.g., 11), computational modeling of lesioned memory performance (12), experimental manipulations of sleep (for review, see 13), pharmacological manipulations of specific neurotransmitter agonists and antagonists (e.g., 14,15), as well as studies of neural replay in animals (16) and humans (17). Despite this broad interest within the domains of neuroscience and computational modeling, there have been relatively few attempts to manipulate consolidation processes within the span of a behavioral laboratory experiment. Hence, many cognitive questions about consolidation processes remain unaddressed. For instance, does consolidation require supra-threshold reactivation of a memory trace? Is consolidation influenced by motivational states or intentionality? Does consolidation demand attentional capacity?

In pursuit of such cognitive questions, researchers have more recently developed an intriguing experimental paradigm using “wakeful rest,” a brief period (roughly 10 minutes) of minimal stimulation while individuals are awake. In a typical wakeful rest paradigm, participants encode memory targets, then often complete an expected *immediate* memory retrieval test for those targets. Experimenters then manipulate the intervening period to include either (a) wakeful rest, which takes place alone in a quiet, comfortable room, without access to electronic devices or reading materials, or (b) a simple distractor task, such as a spot-the-difference picture game (e.g., 18). Finally, participants complete a surprise *delayed* retrieval test for all previous memory targets. The typical finding is that performance on the delayed retrieval task is better for materials that precede wakeful rest than for those that precede the distractor task. We will henceforth refer to this general behavioral pattern as the “wakeful rest effect.”

The wakeful rest effect has been demonstrated using a variety of target stimuli, including prose stories (18–20), word lists (21,22, Experiment 1), Icelandic-English word pairs (23), pronounceable nonwords (22, Experiment 2), virtual maps (24–26), and photographs of everyday objects (27). In addition, a wide variety of distractor tasks have been employed, including spot-the-difference games, visual search tasks, cued autobiographical recall, and passive listening tasks. Importantly, these distractor tasks often overlap minimally with the target stimuli in terms of semantic content and stimulus domain. Thus, one would expect these distractor materials to produce only minimal interference with the target materials. However, wakeful rest effects have been robust, even for these minimally-overlapping materials.

Despite the multiple replications of the wakeful rest effect across a variety of stimulus types and distractor tasks, there have been some recent studies that have failed to observe the effect. Specifically, Varma and colleagues (28) failed to detect a memorial benefit of wakeful rest in comparison to an n-back distractor task, which placed strong cognitive demands on executive resources, but did not overlap with memory targets in semantic or autobiographical processing. Across six experiments, this null effect was consistent for recognition of incidentally-encoded picture-word pairs or faces, as well as free recall of intentionally-encoded auditory words. Moreover, the null effect was consistent using a standard 2-back distractor task, a more engaging, dynamically-adjusting 2-back task, as well as a 3-back task. Varma and colleagues (28) concluded that, since the n-back task did not demand hippocampal-dependent autobiographical processing, such as other common distractors used in the wakeful rest paradigm, it did not interfere with consolidation processes. Additionally, Martini and colleagues (29) failed to demonstrate a wakeful rest effect in two experiments that used a recall task in the participants’ first language (i.e., German) for prose passages studied in a second proficient language (i.e., English), suggesting that the wakeful rest effect might not be generalizable to a wide range of testing conditions. Given these recent failures to extend the wakeful rest effect, it is important to consider to what extent the effect is robust and generalizable.

In addition to potential concerns about replication and generalizability, there are also unresolved questions regarding the theoretical interpretation of wakeful rest memory benefits. Researchers have typically interpreted wakeful rest effects to be a consequence of memory consolidation processes, arguing that, like sleep, wakeful rest is a state that allows for the spontaneous neuronal reactivation, reorganization, and strengthening of a recent memory trace (22). One potential alternative interpretation is that the effect is a consequence of rehearsal, or active maintenance of a recent memory trace within a short-term working memory store. Certainly, an unfilled interval after encoding, such as wakeful rest, would afford opportunities to engage in rehearsal, as compared to an intervening distractor task that demands the allocation of executive or attentional capacity. Further, because wakeful rest studies often include an immediate retrieval test (before the wakeful rest or distractor period), participants might expect another upcoming memory test, which would encourage some form of rehearsal or possibly continue to search for the list items that may not have recalled on the immediate test. Under this interpretation, the wakeful rest paradigm might not capture a pure estimate of consolidation processes, but rather might be subject to the influence of attentionally-controlled rehearsal.

Indeed, previous researchers have been aware of this potential confound and have taken steps to rule out rehearsal as a possible mechanism producing the effect. For example, most studies have included a post-experiment questionnaire to identify participants who expected the surprise memory test or rehearsed targets during the intervening periods. Interestingly, the wakeful rest effect has been consistent whether or not those participants were included in the analyses (21,22). However, since the questionnaires were limited to retrospective self-reports, participants might have been biased to under-report expectation of the test or rehearsal attempts due to demand characteristics. Further, since the questionnaires were typically given at the end of the experiment, it is possible that participants might not have accurately remembered their experiences from earlier in the experiment. Beyond the questionnaire, another approach to minimize rehearsal has been to use pronounceable nonwords as targets. Specifically, Dewar and colleagues (22, Experiment 2), presented nonwords auditorily under the guise of foreign names. The authors argued that since these nonwords produced 0% recall accuracy in separate pilot testing, it was impossible for participants to rehearse them, and yet participants still exhibited a wakeful rest memory benefit for these items in delayed recognition tests. However, since the nonwords were verbally pronounceable, they could be assigned verbal labels, thus potentiating rehearsal. Although admittedly performance would be low for nonwords, it is possible that participants are able to rehearse at least short lists of nonwords (see 30,31). Further, the encoding conditions of this experiment might have encouraged participants to attempt rehearsal. Specifically, the nonwords were presented with explicit encoding instructions to remember the names for a future test. Unlike other similar paradigms, encoding in this experiment was not followed by an immediate memory test, which presumably would serve to limit expectation of the delayed memory test, but see additional concerns above about the influence of an immediate test. In any case, the delayed retrieval test was *not* a surprise to participants, which might have motivated them to engage in rehearsal during the intervening period.

In addition to considering the competing theoretical frameworks of consolidation and rehearsal, one other important question for the wakeful rest paradigm involves the influence of aging. This factor is of particular interest because the study of age-related effects on episodic memory has typically focused on age deficits in encoding and retrieval processes (for review, see 32,33). In contrast, age differences in consolidation are relatively underexplored. Hence, if it is truly representative of consolidation processes, the wakeful rest paradigm might provide novel insight regarding potential age differences in this stage of memory processing. Craig and colleagues (25) recently demonstrated that the wakeful rest benefit for cognitive map accuracy was comparable for both younger and older adults. This observation might suggest that consolidation processes, as measured in the wakeful rest paradigm, are relatively preserved in aging, but further work is needed to replicate this finding and generalize it to other types of memory targets and tests.

In order to further explore the nature of the mechanisms underlying the wakeful rest effect, we conducted a series of experiments that compared both rehearsable and non-reheasable list items, under incidental learning conditions in both younger and older adults. Figure 1 presents the general design of the experiments. In our first experiment, we attempt to replicate the wakeful rest effect in a modified task, which limited the possibility of rehearsal. First, participants encoded memory targets via incidental pleasantness ratings, as opposed to intentional encoding instructions. Incidental encoding should be less likely to motivate rehearsal or retrieval practice during the intervening periods. Additionally, we used abstract shapes as memory targets, which in contrast to prose passages, word lists, nonwords, virtual maps, and familiar objects, are less likely to be rehearsed, as they lack verbal labels to mediate rehearsal. As in previous wakeful rest experiments, each encoding block was followed by either a period of wakeful rest or a distractor task. Finally, memory retention was tested with a surprise retrieval task, followed by a questionnaire to assess whether individuals expected the test and/or attempted to rehearse the targets. By using an incidental encoding paradigm and testing memory for abstract shapes, this study represents, to our knowledge, the strongest attempt to limit rehearsal in the wakeful rest paradigm. However, to anticipate, we did not replicate the expected wakeful rest benefit under these conditions. Across the subsequent experiments, we considered the degree to which the wakeful rest effect might be driven by particular methodological factors that could influence the potential for rehearsal. Specifically, we manipulated both the characteristics of encoding and the distractor tasks. Regarding encoding, we varied whether the task was intentional (Experiment 2) or incidental (all other experiments), and also varied the types of the memory targets, including abstract shapes (Experiments 1-3), nonwords (Experiments 1 and 2), or word lists (Experiments 4-6). Regarding retrieval, we tested for wakeful rest effects in both recognition (all experiments) and free recall tasks (Experiments 4-6). Moreover, we included distractor tasks that involved autobiographical cuing (Experiments 1 and 2), digit monitoring (Experiments 3, 4, and 6), or visual comparison (Experiment 5). Finally, we tested to what extent the wakeful rest effect differs across age groups, by comparing younger and older adult samples (Experiments 3, 4, and 5).

**Fig 1.**
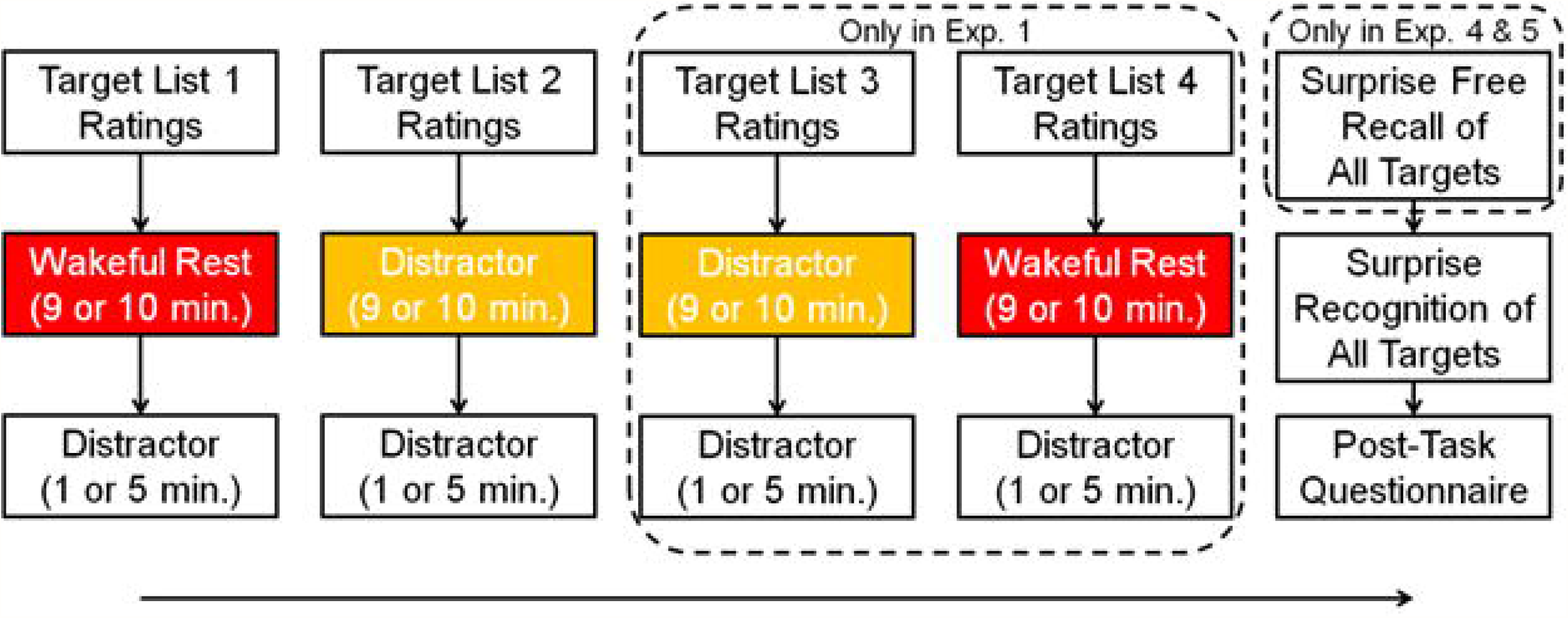
**Basic procedure for Experiments 1, 3, 4, and 5**.

## Experiment 1

Experiment 1 served as a pilot study to test whether the wakeful rest effect replicates under conditions designed to maximally limit rehearsal, including an incidental encoding paradigm and abstract shape targets.

### Method

#### Participants

In all experiments, we used strict screening criteria to eliminate participants who expected the surprise memory test, attempted to rehearse memory targets, or fell asleep, as determined via self-report on a post-task questionnaire. These participants were replaced by new participants in the same counterbalance configuration until the final sample was met. In all experiments, we report only analyses on the final sample, excluding the replaced participants. Importantly, for all analyses, the interpretations do not change if we use a more liberal criteria including all participants.

Experiment 1 participants included a final sample of 16 younger adults recruited from the undergraduate participant pool at Washington University in St. Louis. All participants completed the experiment in exchange for course credit.

#### Design

Experiment 1 included four incidental encoding lists (2 shape and 2 nonword lists) and a surprise final recognition memory test (see Fig 1). Each incidental encoding list was immediately followed by a 9-minute intervening period and then a 1-minute distractor task. The within-participant factors included the type of targets presented at encoding (either non-words or abstract shapes) and the activity that filled the post-encoding period (either wakeful rest or a distractor task).

#### Materials

Sixty nonwords were selected from the English Lexicon Project (34) to use as stimuli. All nonwords were pronounceable combinations of syllables, but were not English words. These nonwords were divided into four lists of 15 nonwords. Lists were matched in the distributions of initial letters, total number of letters, and number of syllables, such that each nonword had a similar lure in each of the other lists. For example, the similar nonwords “accition”, “achuring”, “adduring”, and “afferech” each appeared in separate lists. Nonwords were presented visually in black 24-point Arial font on a white background.

We also selected 60 abstract shapes, randomly generated and normed by Vanderplas and Garvin (35). These shapes were divided into four lists of 15 shapes. Lists were matched in the distributions of total number of points and association value (i.e., the degree to which the shape reminded participants of something meaningful, according to Vanderplas & Garvin, 1951), such that each shape had a similar lure in each of the other lists. Shapes were presented filled in with black on a white background.

All stimuli were presented using E-Prime 2.0 software (Psychology Software Tools, Pittsburgh, PA). For each participant, two lists of nonwords and two lists of shapes were selected as targets to be presented at the various encoding phases, while the other two served as lures at the recognition test. Across participants, each list appeared an equal number of times as a target or as a lure, an equal number of times in the wakeful rest condition or in the distractor condition, and an equal number of times in each serial encoding position, i.e., first, second, third, or fourth.

#### Procedure

All testing took place in a dimly lit room with minimal furniture and no reading materials. Participants did not have access to phones or any other personal belongings for the duration of the experiment. Participants first completed a practice round of the digit monitoring task. In this task, participants listened to audio recordings of digits, ranging from 0 to 9, presented in a pseudorandom order at a rate of one every three seconds. Participants were instructed to press the space bar when they heard an odd digit that was preceded by another odd digit, which occurred on 20% of the trials. Following the instructions and practice with the distraction task, participants completed four rounds of incidental encoding: two rounds for nonword targets and two rounds for shape targets. Participants were instructed through a false cover story that the experiment primarily addressed how different types of stimuli are rated for pleasantness. They were not informed that they would complete any memory tests. In each round, participants viewed 15 target items for three seconds each. They were instructed to rate the pleasantness of each item via button press on a Likert scale from 1 (for very unpleasant) to 5 (for very pleasant). After each round of pleasantness ratings, the experimenter provided a false cover story that the targets in the next round would be selected after the experimenter analyzed the previous pleasantness ratings, thus requiring a break of approximately 9 minutes. This interval was filled by either wakeful rest or a distractor task, the sequence of which was counterbalanced across participants.

During the 9-minute wakeful rest interval, participants were left alone in the testing room. The testing computer was password-locked with a blank monitor, so as not to distract them. Participants were instructed to rest quietly and to try not to fall asleep.

During the 9-minute distractor interval, participants were left alone in the testing room. The testing computer was password-locked with a blank monitor. At pseudo-random intervals of on average 46 seconds, the computer played an audio recording of a recognizable event, e.g., raindrops falling. Following the paradigm used by Craig and colleagues (21, Experiments 3 and 4), participants were instructed to rest quietly and that the recordings would be presented in order to prevent them from falling asleep.

After each 9-minute interval, participants completed a 1-minute round of the digit monitoring task as an additional distractor. This task served to minimize any carry-over effects of fatigue on the next encoding list, which may differ between the wakeful rest and distractor conditions, as done in some previous wakeful rest experiments (e.g., 22, Experiment 1). Of course, any benefit of wakeful rest on memory consolidation should be present after this 1 minute washout period.

After completing four cycles of incidental encoding, intervening period, and short distractor tasks, participants completed a final surprise recognition memory test for nonwords and shapes randomly intermixed, including targets and foils. Targets included all nonword and shape lists presented across the four rounds of pleasantness ratings. Foils included the four stimulus lists (two lists of nonwords and two lists of shapes) held in reserve. For each of the 60 items, participants were instructed to indicate by button press whether it had been presented in any of the pleasantness rating tasks (i.e., it was “old”) or it was new. After each memory judgment, they provided a rating of confidence on a Likert scale from 1 (completely guessing) to 5 (completely confident). Participants had no time limit to provide memory judgments or confidence ratings, but were instructed to do so as quickly and as accurately as possible.

Finally, participants completed a post-task questionnaire. Participants were replaced if their questionnaire responses indicated that they (1) had expected a memory test, (2) attempted to rehearse the shapes or non-words, or (3) fell asleep at any point in the experiment.

### Results

#### Questionnaire

Three participants reported expecting the memory test. Two participants reported attempting to rehearse the targets. Each of these participants was replaced by a new participant given the same counterbalancing order of conditions. An additional participant was replaced because it was revealed that they had a smartphone in their possession during the breaks, against instructions.

#### Recognition

Figure 2 displays recognition memory performance (defined as hits – false alarms) as a function of target type and distractor condition. We tested the effects of target type and condition on recognition performance in a 2 x 2 Analysis of Variance (ANOVA), with hits – false alarms as the dependent variable, and target (nonwords or shapes) and condition (wakeful rest or listening) as within-participant factors. This analysis revealed a significant main effect of target type, *F*(1,15) = 26.09, *p* < .001, *η*_*G*_^2^ = .41, but no hint of a main effect of condition, *F*(1,15) = 0.55, *p* = .469, *η*_*G*_^2^ < .01, nor an interaction between target type (nonwords vs. shapes) and condition, *F*(1,15) = 0.23, *p* = .640, *η*_*G*_^2^ < .01. As shown in Figure 2, recognition performance was lower for shapes than it was for nonwords, and performance for both target types was similar in the wakeful rest condition, as compared to the listening condition.

**Fig 2.**
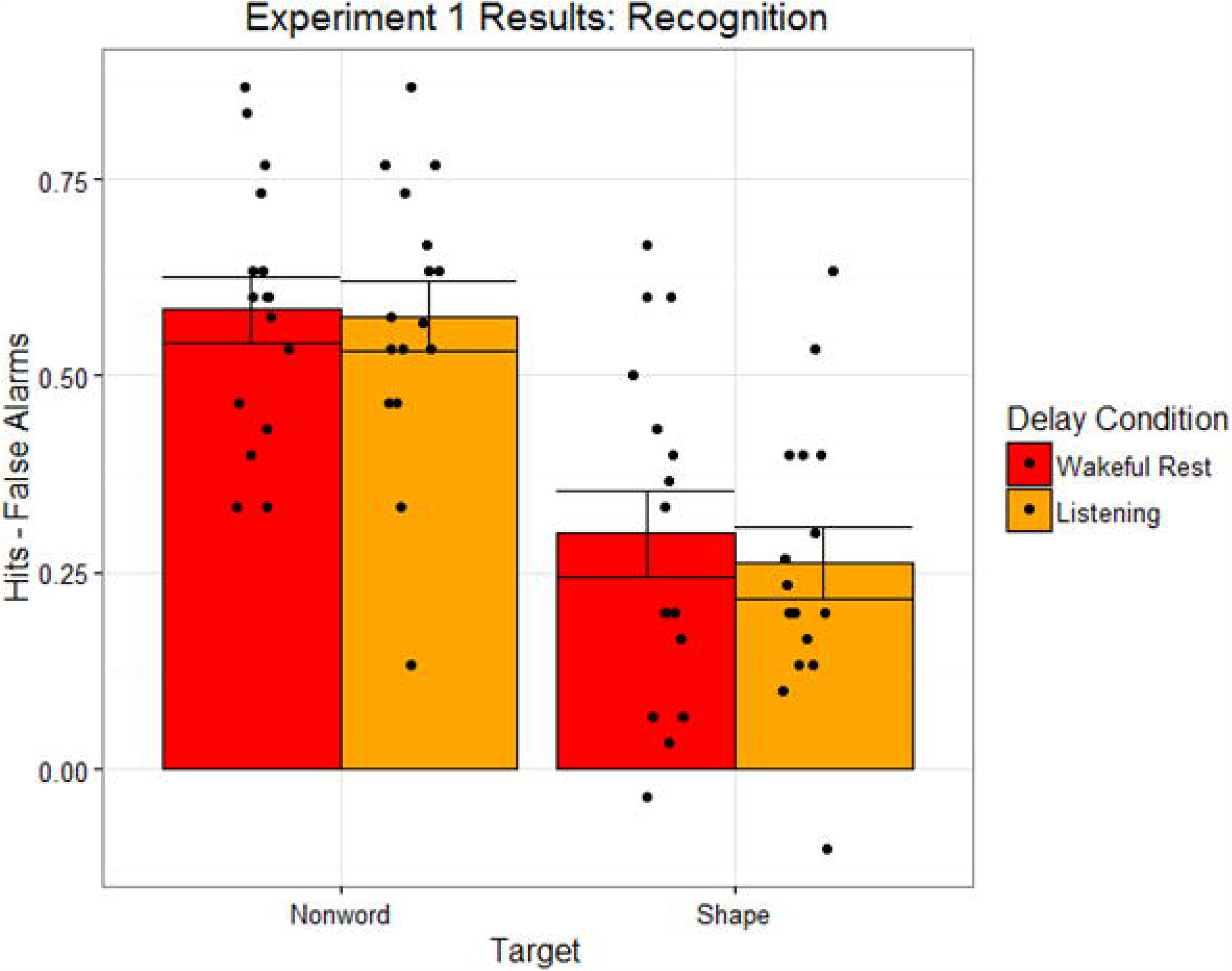
Recognition task performance for nonword and shape targets, as a function of distractor condition, in Experiment 1. Bar heights reflect mean values. Points reflect individual participants. Error bars are standard error of the mean.

The null effect of condition on recognition accuracy is inconsistent with the several earlier demonstrations of the wakeful rest benefit for various memory tests (see Introduction), but is consistent with recent failures to obtain the effect. However, the interpretation of this null result is obviously limited by the possibility of insensitivity in the current dataset. Recently, Bayes factors have been proposed as a framework to arbitrate whether observed data in a null effect support a null hypothesis or are insensitive to a possible alternative hypothesis (36). To address this concern, we reanalyzed the data using a repeated-measures Bayesian ANOVA approach, implemented using the ‘BayesFactor’ package in R (37), using the default prior values, as recommended by Rouder and colleagues (38). Critically, for the effect of condition, the estimated Bayes factor, as compared against a null model (*BF*_*10*_), was 0.277. Hence, the data shift the relative plausibility of an alternative hypothesis over the null by only a factor of 0.277 (or, correspondingly, the data support the *null* over an *alternative* hypothesis, *BF*_*01*_, by a factor of 3.606. This value implies that the data provide substantial evidence in favor of the null hypothesis (see 39).

### Discussion

The results from Experiment 1 indicate that memory for nonwords and abstract shapes does not benefit from a minimal post-encoding period of wakeful rest. This result is inconsistent with a previous demonstration of a wakeful rest effect for intentionally-encoded nonword stimuli (22). However, there are a number of plausible reasons why we did not find an effect in this study, which will be further considered across the subsequent experiments.

First, memory targets were incidentally encoded in the current experiment, but were intentionally encoded in most prior demonstrations. As a reminder, in previous studies, participants intentionally encoded targets, usually aware of at least an immediate memory test before the intervening task, but presumably did not anticipate a surprise delayed test (18,e.g., 21,22). As noted earlier, we believe that the mere presence of the immediate memory test might increase the likelihood of participants thinking about the list items during the subsequent wakeful reset period.

Second, the current distractor task requires minimal engagement and differs only slightly from the wakeful rest condition. Thus, this manipulation of the intervening period might not be sufficiently strong to modulate consolidation processes, although it should be noted that the wakeful rest effect has been demonstrated for recall of intentionally-encoded words using a similar manipulation of distractor condition (21, Experiments 3 and 4).

Third, the current methods differed from previous demonstrations of the effect in that all variables were manipulated within participants and memory was tested with a recognition task. Indeed, the wakeful rest effect has been demonstrated in experiments using a within-participant design (18,e.g., 21) and in experiments using a recognition task (22,e.g., 27), but to our knowledge, never in an experiment using both elements.

Fourth, considering the small sample size of this pilot study (N = 16), there might be inadequate power to detect an effect of the relatively weak manipulation, but given the substantial evidence in favor of the null offered by the Bayes factor analyses, we decided to examine alternative constraints on the effect.

Fifth, memory target stimuli differed from previous demonstrations of the effect. Nonwords have been used as targets in one prior experiment (22, Experiment 2), but they were presented auditorily, disguised as foreign names, in contrast to the visual presentation in the current experiment. Importantly, the nonwords were presented under intentional learning conditions. Abstract shapes, as mentioned, have never been used in any wakeful rest experiment.

## Experiment 2

In Experiment 2, we tested whether nonword and abstract shape stimuli, similar to those used in Experiment 1, would experience a wakeful rest benefit when they are encoded intentionally for an immediate memory test. We also substantially increased the sample size in this and subsequent experiments in order to increase statistical power over the pilot, Experiment 1.

### Method

#### Participants

Experiment 2 participants included a final sample of 32 younger adults recruited from the undergraduate participant pool at Washington University in St. Louis. All participants completed the experiment in exchange for course credit.

#### Design

The design was similar to the one described in Experiment 1 with one major distinction: encoding was intentional instead of incidental. In order to minimize expectation of the final *delayed* memory test in the intentional encoding paradigm, each encoding phase was followed by an immediate recognition task, before the wakeful rest or distractor period (see Fig 3). A similar immediate recall procedure has been employed in several demonstrations of the wakeful rest effect, in order to assess memory retention from encoding to after the intervening period (18,e.g., 21,22, Experiment 2,23). This design presumably limits participants’ expectation of the final surprise memory test in an intentional encoding paradigm, since from the participant’s perspective they have already been tested for the lists. As in Experiment 1, we manipulated, within participants, the type of targets presented at encoding (either non-words or abstract shapes) and the activity that filled the post-encoding interval (either wakeful rest or a distractor task).

**Fig 3.**
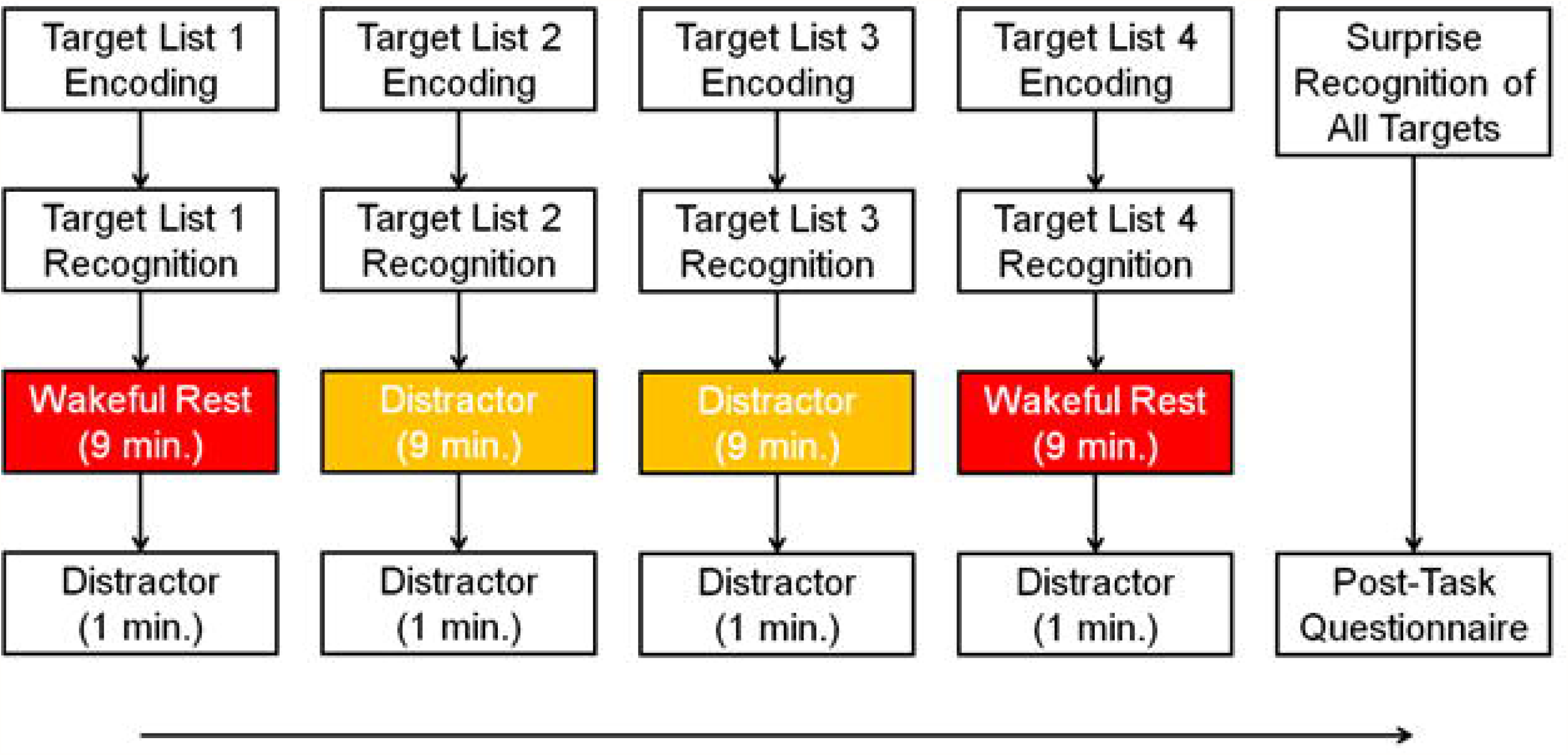
**Basic procedure for Experiment 2**.

#### Materials

Stimuli were drawn from the same source materials as described in Experiment 1. In Experiment 2, we used a greater number of stimuli per list: including four lists of 24 nonwords and four lists of 24 shapes. Lists were constructed based on the same controls and were rotated with the same counterbalancing scheme described in Experiment 1.

#### Procedure

The procedure was similar to the one described in Experiment 1. Aspects of the procedure were slightly changed to allow for intentional encoding instructions and immediate recognition memory tests following each encoding phase, as described below.

Before each encoding phase, participants were informed that their memory for the targets would be tested in an immediate recognition test, as in previous wakeful rest experiments (see 18, etc.). They were instructed to study each target and provide a pleasantness rating from 1 to 5 for each item within a 3-second time limit.

After each encoding phase, participants completed an immediate recognition memory test for the previously-studied items. Targets included 6 of the 24 items in the previously-encoded list. Foils included 6 of the 24 items from a list held in reserve. Participants provided confidence ratings from 1 (completely guessing) to 5 (completely confident) for each memory judgment. Participants had no time limit to provide memory judgments or confidence ratings, but were instructed to do so as quickly and as accurately as possible.

After immediate recognition, participants were instructed that the next memory target list would be selected based on their previous pleasantness ratings and recognition performance, thus requiring a break of approximately 9 minutes. This interval was filled by either wakeful rest or a distractor task. The wakeful rest phase was exactly the same as the procedure described in Experiment 1, but the listening distractor phase was slightly different. Following the paradigm used by Craig and colleagues (21, Experiments 1 and 2), for each recording, participants were instructed to imagine a specific event related to that sound, either from their past or in their future. For example, if presented with a cat’s meow, they might think of an interaction with a pet. They were instructed to think about the event in as much detail as possible for as long as possible, until the next recording was played. Participants were given writing materials and were instructed to write down details describing their imagined events. Hence, we anticipated that this manipulation would produce more disruption during the distractor phase compared to the passive listening condition used in Experiment 1.

In the final surprise recognition task, participants discriminated between 72 targets, composed of the 18 items from each of the 4 studied lists that were not presented in the immediate tests, and 72 lures, composed of the 18 items from each of the 4 reserve lists that were not presented as lures in the immediate tests. Apart from the differences described here, the procedure was identical to the one described in Experiment 1.

### Results

#### Questionnaire

Twenty-four participants (38%) reported expecting the final memory test. Thus, as we suspected, when given intentional encoding instructions and an immediate test there is an increase in the likelihood of expecting a final test, at least compared with Experiment 1 (14%). Seven participants reported attempting to rehearse the targets. Two participants reported falling asleep. Each of these participants was replaced by a new participant given the same counterbalancing order of conditions.

#### Immediate Recognition

We first examined performance in the immediate recognition test, to validate that any differences in the delayed test were due to the manipulation of the intervening period, and not any difference in initial encoding strength. Figure 4 displays recognition memory performance (defined as hits-false alarms) on the immediate tests as a function of target type and distractor condition. We tested the effects of target type and condition on immediate recognition performance in a 2 x 2 ANOVA, with hits-false alarms as the dependent variable and with target (nonwords or shapes) and condition (wakeful rest or listening) as within-participant factors. This analysis revealed a significant main effect of target type, *F*(1,31) = 52.26, *p* < .001, *η*_*G*_^2^ = .30, but not a main effect of condition, *F*(1,31) = 0.72, *p* = .404, *η*_*G*_^2^ < .01, nor an interaction between target and condition, *F*(1,31) = 1.31, *p* = .261, *η*_*G*_^2^ = .01. As shown in Fig 4, immediate recognition performance was again lower for shapes than it was for nonwords, and performance for both target types was similar in the wakeful rest condition, as compared to the listening condition.

**Fig 4.**
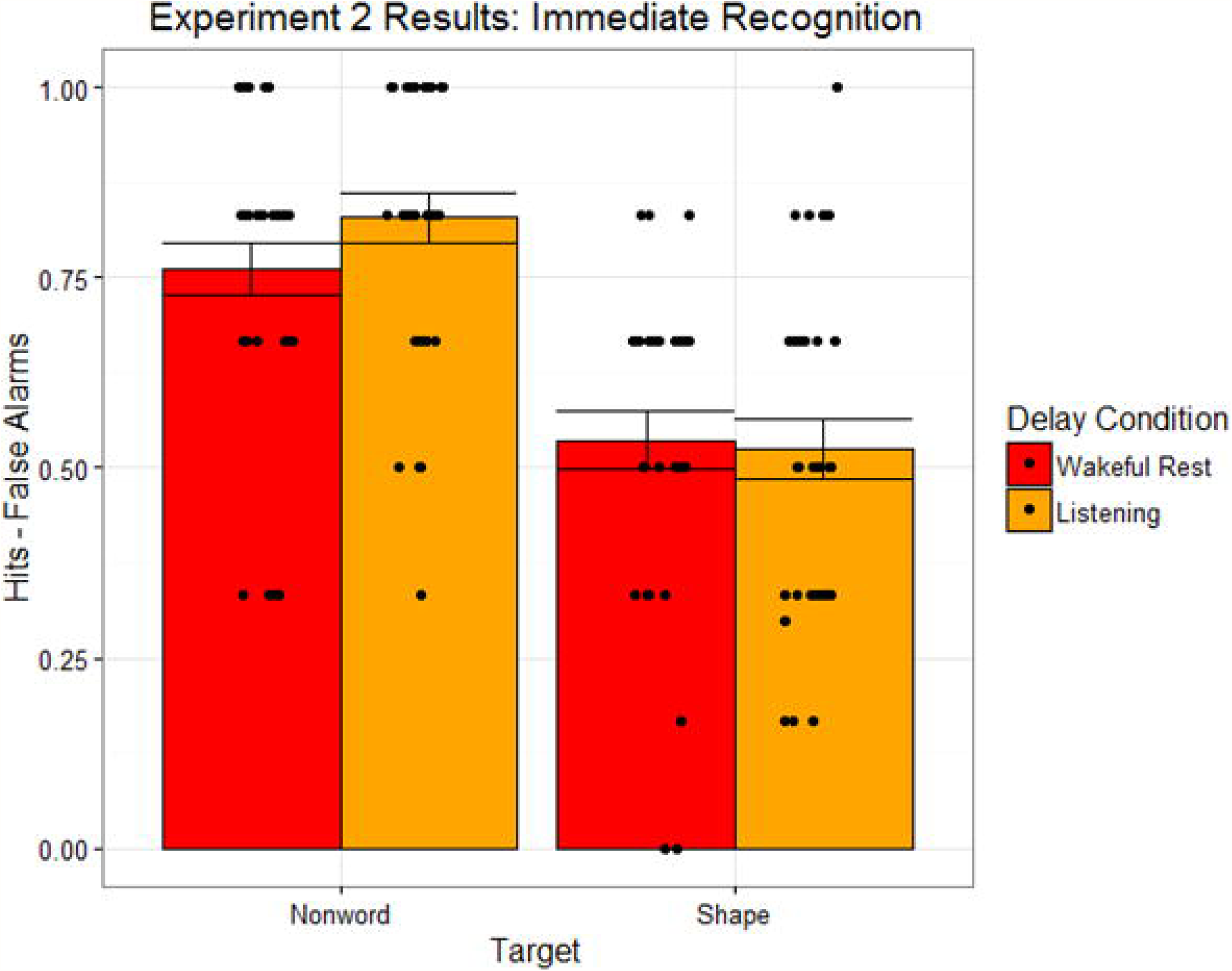
Immediate recognition task performance for nonword and shape targets, as a function of distractor condition, in Experiment 2. Bar heights reflect mean values. Points reflect individual participants. Error bars are standard error of the mean.

#### Delayed Recognition

Figure 5 displays final recognition memory performance (defined as hits-false alarms) on the delayed test as a function of target type and distractor condition. We tested the effects of target type and condition on delayed recognition performance in a 2 x 2 ANOVA, using the same factor structure described for immediate recognition. This analysis revealed a significant main effect of stimulus type, *F*(1,31) = 26.95, *p* < .001, *η*_*G*_^2^ = .24, but not a main effect of condition, *F*(1,31) = 0.07, *p* = .787, *η*_*G*_^2^ < .01, nor an interaction between target and condition, *F*(1,31) = 0.35, *p* = .557, *η*_*G*_^2^ < .01. We followed up on the null effect of condition with a Bayesian repeated-measures ANOVA. The estimated Bayes factor for the effect of condition, *BF*_*10*_, was 0.188. Hence, the data provide substantial support for the null over an alternative hypothesis, *BF*_*01*_, by a factor of 5.319. As shown in Fig 5, delayed recognition performance was lower for shapes than it was for nonwords, and performance for both target types was similar in the wakeful rest condition, as compared to the listening condition.

**Fig 5.**
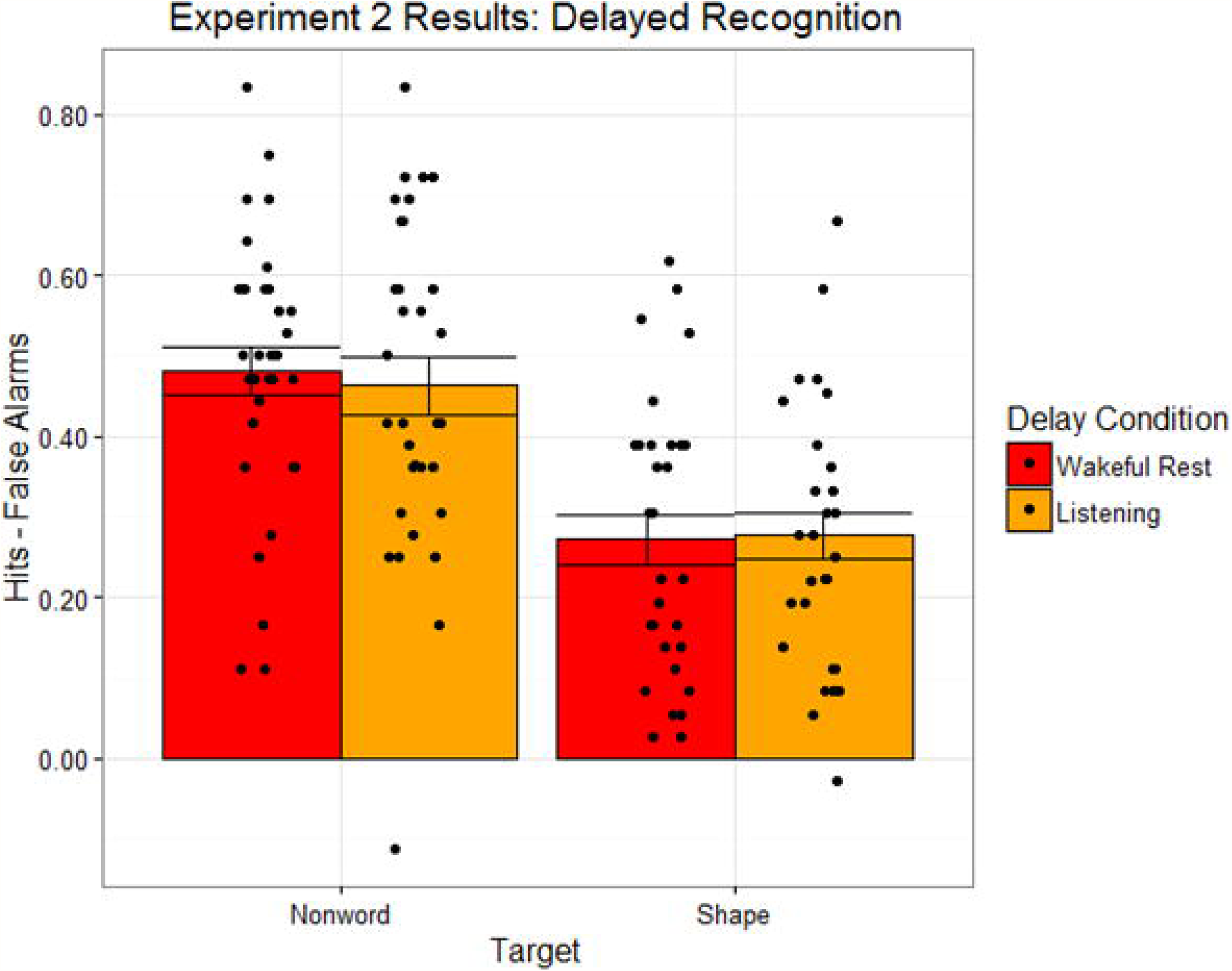
Delayed recognition task performance for nonword and shape targets, as a function of distractor condition, in Experiment 2. Bar heights reflect mean values. Points reflect individual participants. Error bars are standard error of the mean.

#### Retention

Figure 6 displays the difference in recognition performance (defined as hits-false alarms) between the immediate and delayed recognition tests as a function of target type and distractor condition to examine if there are retention differences as a function of condition. Values are plotted such that negative values indicate lower performance on the delayed test than on the immediate test, or lower retention. We tested the effects of target type and condition on retention in a 2 x 2 ANOVA, using the same factor structure described for immediate recognition. This analysis revealed no significant main effects of target type, *F*(1,31) = 1.44, *p* = .240, *η*_*G*_^2^ = .02, condition, *F*(1,31) = 0.89, *p* = .352, *η*_*G*_^2^ = .01, nor an interaction between target and condition, *F*(1,31) = 2.88, *p* = .100, *η*_*G*_^2^ = .01. Again, we followed up on the null effect of condition with a Bayesian repeated-measures ANOVA. The estimated Bayes factor for the effect of condition, *BF*_*10*_, was 0.274. Hence, the data provide substantial support for the null over an alternative hypothesis, *BF*_*01*_, by a factor of 3.650. As shown in Fig 6, retention was similar for shapes and nonwords, and retention for both target types was similar in the wakeful rest condition, as compared to the listening condition.

**Fig 6.**
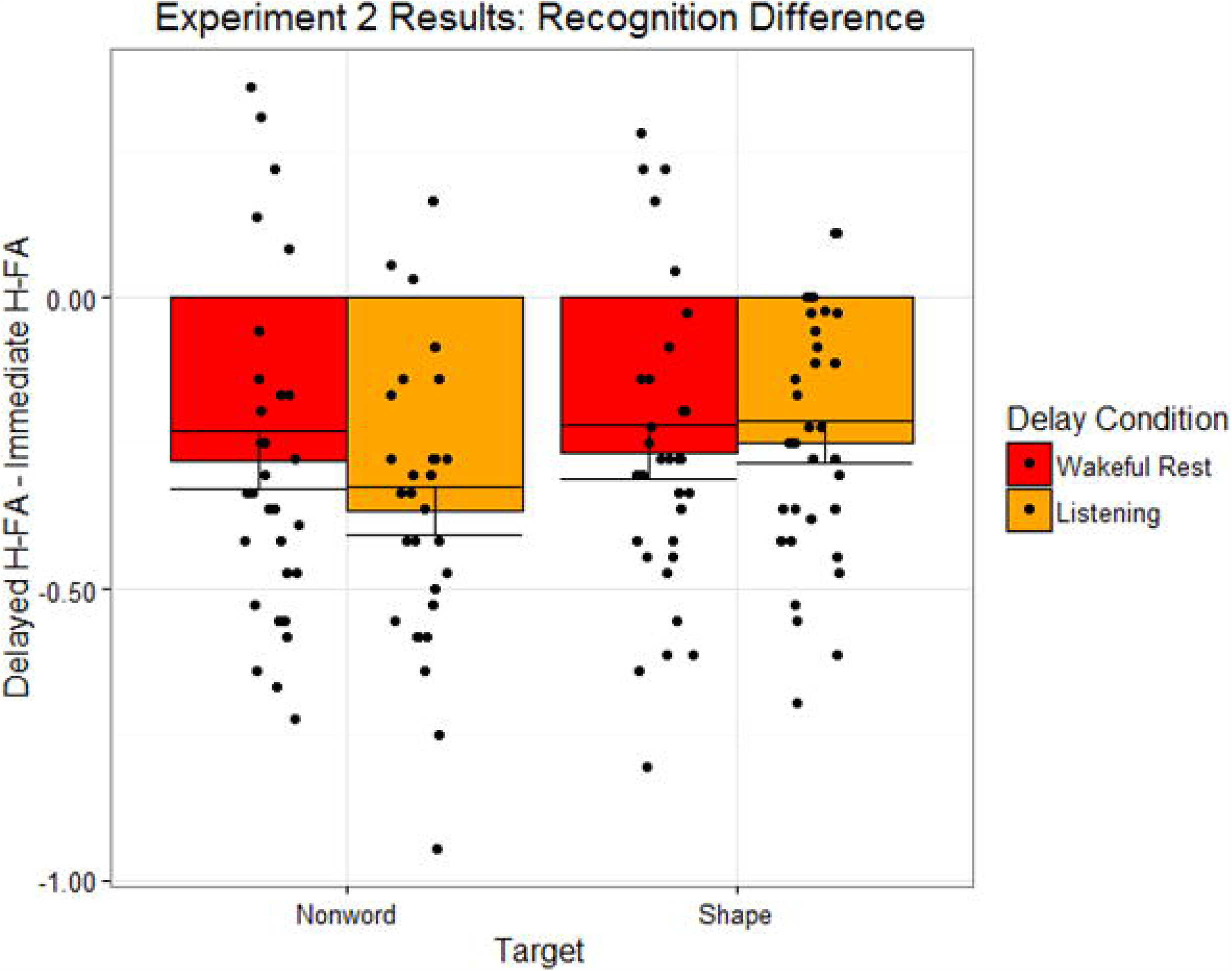
Recognition task performance difference for nonword and shape targets, as a function of distractor condition, in Experiment 2. Bar heights reflect mean values. Points reflect individual participants. Error bars are standard error of the mean.

### Discussion

The results from Experiment 2 demonstrate that, using intentional encoding instructions, memory for nonwords and abstract shapes still does not benefit from a post-encoding period of wakeful rest. This pattern is inconsistent with a previous demonstration of a wakeful rest effect for intentionally-encoded nonword stimuli (22). Of course, it is still possible that autobiographical listening is a relatively weak manipulation of the post-encoding interval, even though a similar procedure produced a wakeful rest effect for intentionally encoded words (21), and indeed this condition has recently been argued to be particularly important distractor task to observe a wakeful reset effect (see 40). It is also possible that mixing the nonwords with the abstract patterns may have minimized the wakeful rest effect for some unknown reason. Previous experiments have only included one type of stimulus throughout the experiment.

## Experiment 3

In Experiment 3, we included abstract shapes as the only memory targets. We tested whether these stimuli would exhibit a wakeful rest benefit in comparison to a more demanding digit monitoring distractor task. We eliminated the immediate test condition in this experiment, because it appears to increase the likelihood of rehearsal. Finally, we also included an older adult participant group to test whether the strength of the wakeful rest effect differed between age groups. As mentioned previously, there has been evidence that the wakeful rest effect for verbal materials occurs in both younger adult samples (21,23) and healthy older adult samples (18,22), but to our knowledge, potential age differences in the effect have not been directly tested in the same study.

### Method

#### Participants

Experiment 3 participants included a final sample of 32 younger adults and 32 older adults. Younger adults were recruited from the undergraduate participant pool at Washington University in St. Louis and completed the experiment in exchange for course credit. Older adults were recruited from the Volunteer for Health registry at Washington University in St. Louis and completed the experiment in exchange for cash payment. Older adults were screened for possible dementia using the Short Blessed Test (41). All participants scored within the range of normal cognition (0-4 out of 28, 42), and so no participants were excluded for cognitive impairment.

#### Design

The design was similar to the one described in Experiment 1 with one major distinction: we tested memory for only abstract shapes and not non-words. To that end, we only included two incidental encoding phases (see Fig 1). Within participants, we manipulated the activity that filled the post-encoding interval (either wakeful rest or a distractor task). Age group was a between-participant variable (younger or older adults).

#### Materials

Stimuli included only the abstract shape stimuli described in Experiment 1, consisting of four lists of 15 shapes.

#### Procedure

The procedure was similar to the one described in Experiment 1. Aspects of the procedure were slightly changed to test the effect using a stronger manipulation of the post-encoding interval, as described below.

Experiment 3 included only two incidental encoding phases. During each phase, participants incidentally encoded a list of abstract shapes by providing pleasantness ratings, as described in Experiment 1.

The post-encoding interval lasted 10 minutes. It consisted of either wakeful rest or a 10-minute interval of the digit monitoring task described in Experiment 1. After each 10-minute interval, participants completed the same digit monitoring task for an additional 5 minutes, a longer duration than in Experiments 1 and 2, in order to more effectively eliminate fatigue accounts of any conditional differences, as per Dewar and colleagues (18), etc. As noted above, the purpose of this phase was to minimize any differential carry-over effects of fatigue on subsequent encoding or recognition tasks. Of course, any benefits of the 10 minute wakeful rest should persist across this wash-out period.

### Results

#### Questionnaire

Eight younger and 13 older participants reported some expectancy for a memory test. Four participants (all older) reported attempting to rehearse the targets. Five participants (3 younger and 2 older) reported falling asleep. Each of these participants was replaced by a new participant given the same counterbalancing order of conditions to achieve 32 participants per age group.

#### Recognition

Figure 7 displays recognition memory performance (defined as hits-false alarms) as a function of age and distractor condition. We tested the effects of healthy aging and condition on recognition performance in a 2 x 2 mixed-model ANOVA, with hits-false alarms as the dependent variable, age as a between-participant factor (younger or older adults) and condition as a within-participant factor (wakeful rest or digit monitoring). This analysis revealed a significant main effect of age, *F*(1,62) = 19.00, *p* < .001, *η*_*G*_^2^ = .18, but not a main effect of condition, *F*(1,62) = 0.17, *p* = .680, *η*_*G*_^2^ < .01, nor an interaction between age and condition, *F*(1,62) = 1.07, *p* = .304, *η*_*G*_^2^ = .01. Again, we followed up on the null effect of condition with a Bayesian mixed-model ANOVA. The estimated Bayes factor for the effect of condition, *BF*_*10*_, was 0.200. Hence, the data provide substantial support for the null over an alternative hypothesis, *BF*_*01*_, by a factor of 5.000. As shown in Fig 7, older adults had lower recognition performance than younger adults, and both groups performed similarly in the wakeful rest condition, as compared to the digit monitoring condition.

**Fig 7.**
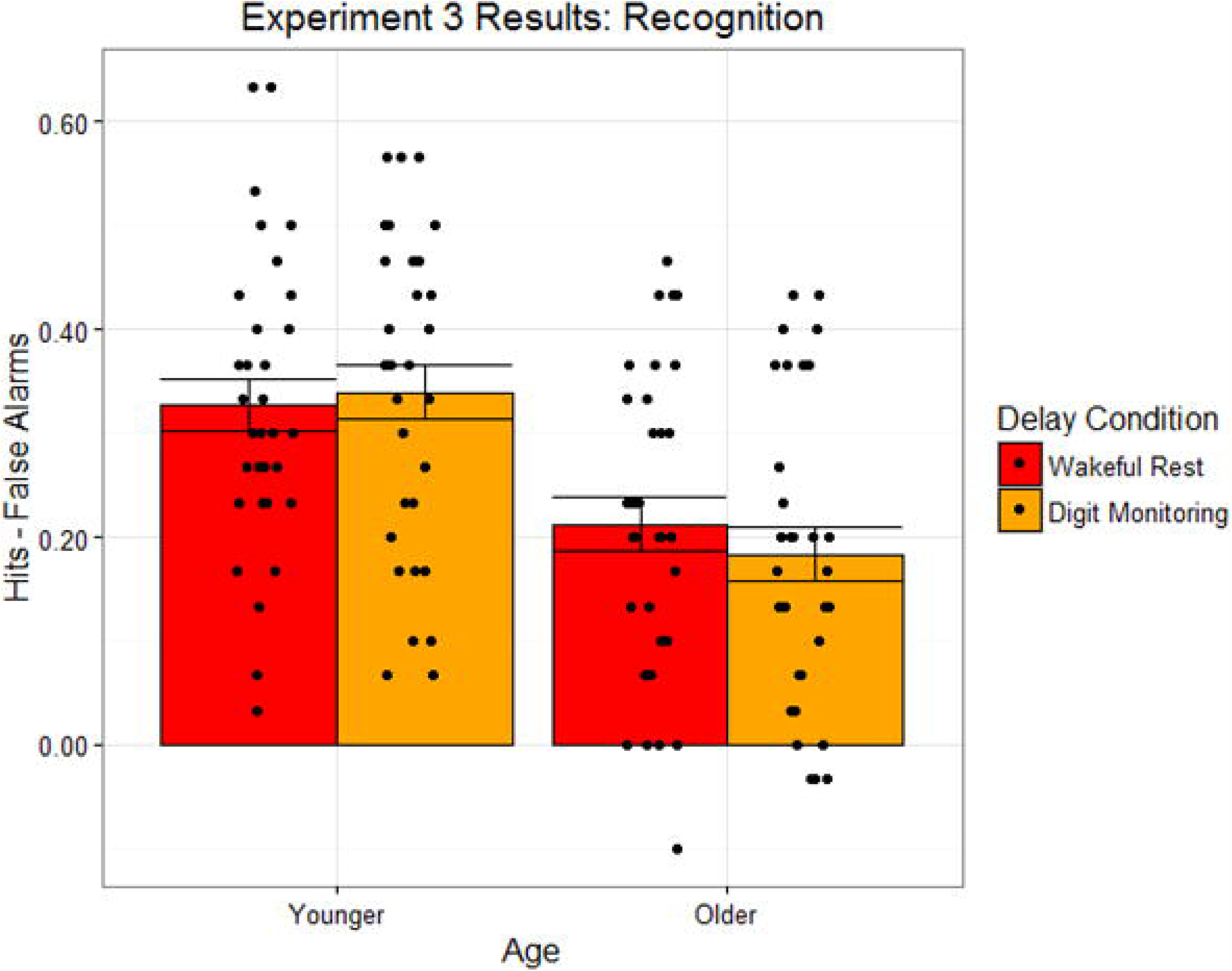
Recognition task performance for shape targets in younger and older adults, as a function of distractor condition, in Experiment 3. Bar heights reflect mean values. Points reflect individual participants. Error bars are standard error of the mean.

### Discussion

The results from Experiment 3 demonstrate that, using a stronger manipulation of the post-encoding interval, memory for abstract shapes still does not benefit from a post-encoding period of wakeful rest. This effect is inconsistent with a previous demonstration of a wakeful rest effect for intentionally-encoded nonword stimuli (22, Experiment 2). It is possible that the shapes presented in the previous experiments are dependent upon a system that is not disrupted by the demanding digit monitoring task, but other types of memory targets, for instance word lists, might be more likely disrupted.

Moreover, it is also possible that the failure to observe the expected wakeful rest effect in Experiments 1 – 3 is related to the use of a recognition memory task. Dewar and colleagues have reported some evidence of wakeful rest effects on recognition task performance (22,27), so the test type should not limit the results. However, recall tasks are more dependent on attentionally-controlled recollection processes than recognition tasks, which may also be influenced by automatic familiarity. Hence, if the wakeful rest effect acts on attentionally-controlled retrieval processes, then recall tasks might be more sensitive to the effect.

## Experiment 4

In Experiment 4, we tested whether different stimuli (i.e., words) would experience a wakeful rest benefit in comparison to the same digit monitoring distractor task used in Experiment 3. In addition, these verbal targets also afford testing memory via a recall task, unlike the prior shapes, which were only tested via recognition in the previous experiments.

### Method

#### Participants

Experiment 4 participants included a final sample of 24 younger adults and 16 older adults, recruited from the same sources described in Experiment 3. Again, on the Short Blessed Test, all older participants scored within the range of normal cognition, and so no participants were excluded for cognitive impairment.

#### Design

The design was similar to the one described in Experiment 3 with one major distinction: we tested memory for visually-presented words, instead of abstract shapes. Within participants, we manipulated the activity that filled the post-encoding interval (either wakeful rest or a distractor task), with age group the only between-participant variable.

#### Materials

Stimuli included 60 unrelated nouns, divided into four lists of 15 words. List words were selected from the English Lexicon Project (34) and matched in the distributions of initial letters, total number of letters, number of syllables, log Hyperspace Analogue to Language (HAL) frequency, orthographic neighborhood, phonological neighborhood, and number of phonemes. Words were presented visually in black 24-point Arial font on a white background.

#### Procedure

The procedure was similar to the one described in Experiment 3. Aspects of the procedure were slightly changed to test the effect in visual word stimuli, as described below.

Experiment 4 included two incidental encoding phases. During each phase, participants incidentally encoded a list of visually-presented nouns by providing pleasantness ratings in a task similar to the ones described in Experiments 1.

As described in Experiment 3, the post-encoding interval lasted 10 minutes. It consisted of either wakeful rest or a 10-minute interval of the digit monitoring task described in Experiment 1. After each 10-minute interval, participants complete the same digit monitoring task for an additional 5 minutes.

After completing both cycles of incidental encoding, intervening period, and short distractor tasks, participants completed a surprise recall memory test. Participants were instructed to vocally recall the words they viewed during the pleasantness rating tasks. An experimenter recorded all responses for at least 60 seconds, and stopped the task when a period of 30 seconds elapsed without a response. After the recall task, participants were given an old-new recognition memory task for previously studied target words and novel lure words.

Apart from the differences described here, the procedure was identical to the one described in Experiment 3.

### Results

#### Questionnaire

Three participants (1 younger and 2 older) reported expecting the memory test. Each of these participants was replaced by a new participant given the same counterbalancing order of conditions.

#### Recall

Figure 8 displays the proportion of words recalled as a function of age and distractor condition. We tested the effects of healthy aging and condition on recall performance in a 2 x 2 mixed-model ANOVA, with proportion recalled as the dependent variable, age as a between-participant factor (younger or older adults) and condition as a within-participant factor (wakeful rest or digit monitoring). This analysis revealed a marginal main effect of age, *F*(1,38) = 3.69, *p* = .062, *η*_*G*_^2^ = .06, a highly significant main effect of condition, *F*(1,38) = 16.89, *p* < .001, *η*_*G*_^2^ = .11, with no interaction between age and condition, *F*(1,38) = 0.24, *p* = .627, *η*_*G*_^2^ < .01. As shown in Fig 8, older adults recalled marginally fewer words than younger adults did, and both groups recalled more words in the wakeful rest condition than in the digit monitoring condition.

**Fig 8.**
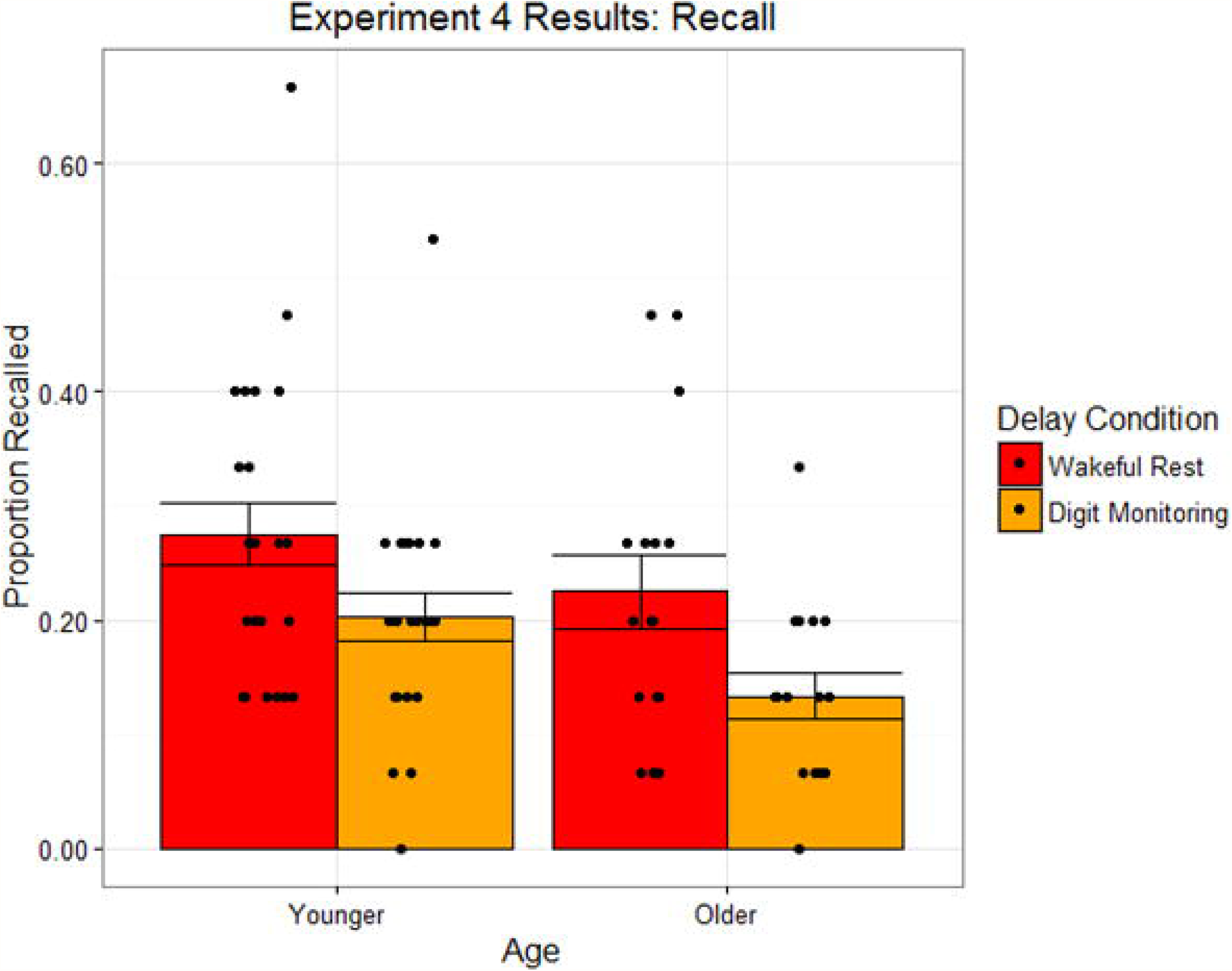
Recall task performance for visual word targets in younger and older adults, as a function of distractor condition, in Experiment 4. Bar heights reflect mean values. Points reflect individual participants. Error bars are standard error of the mean.

#### Recognition

Figure 9 displays recognition memory performance (defined as hits-false alarms) as a function of age and distractor condition. We tested the effects of healthy aging and condition on recognition performance in a 2 x 2 mixed-model ANOVA using the same factor structure described for recall. This analysis revealed no significant main effects of age, *F*(1,38) = 1.46, *p* = .234, *η*_*G*_^2^ < .01, condition, *F*(1,38) = 0.01, *p* = .923, *η*_*G*_^2^ < .01, nor an interaction between age and condition, *F*(1,38) = 0.08, *p* = .783, *η*_*G*_^2^ < .01. Again, we followed up on the null effect of condition with a Bayesian mixed-model ANOVA. The estimated Bayes factor for the effect of condition, *BF*_*10*_, was 0.234. Hence, the data provide substantial support for the null over an alternative hypothesis, *BF*_*01*_, by a factor of 4.274. As shown in Fig 9, older adults had statistically similar, but numerically lower, recognition performance than younger adults did, and both groups performed similarly in the wakeful rest condition, as compared to the digit monitoring condition.

**Fig 9.**
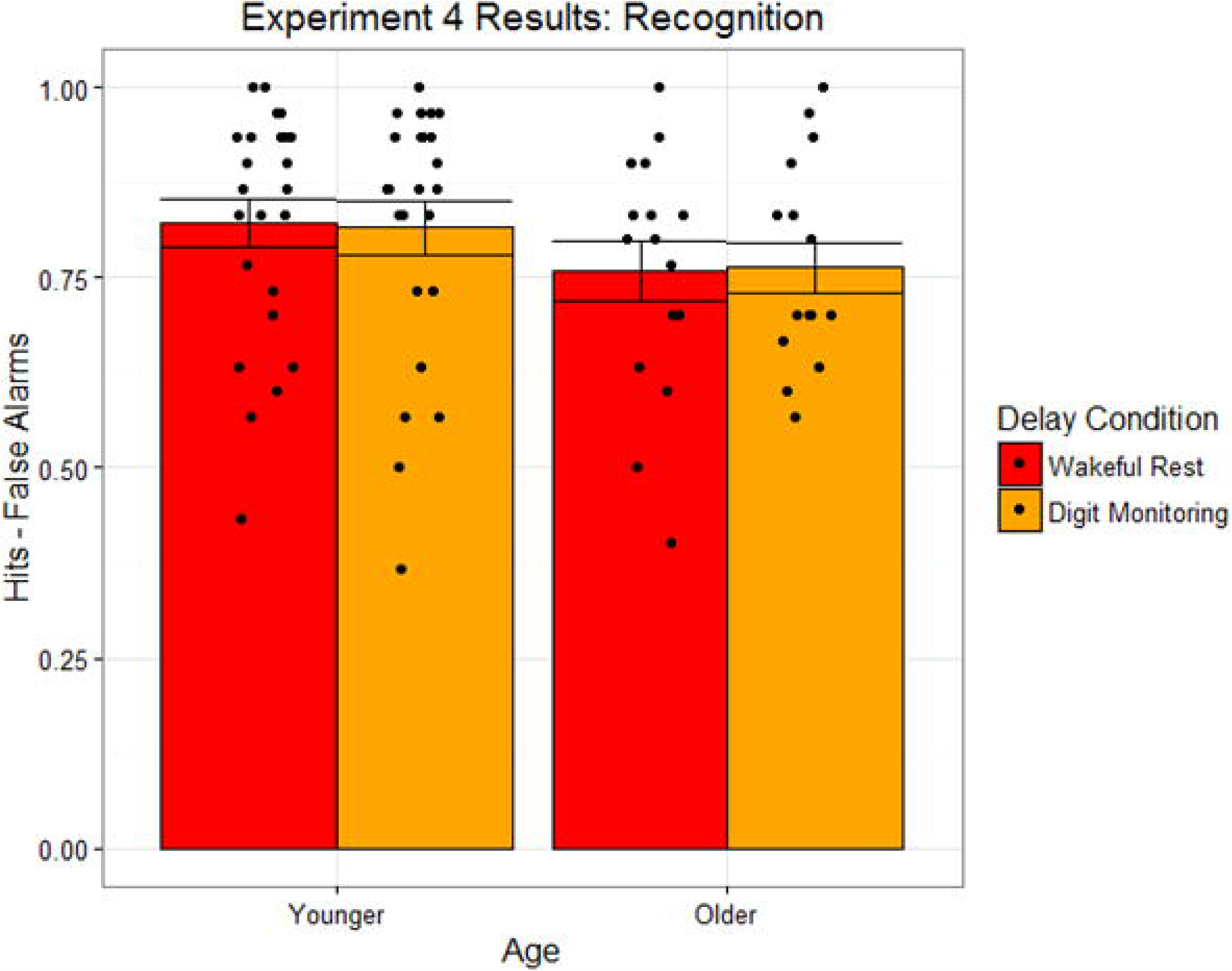
Recognition task performance for visual word targets in younger and older adults, as a function of distractor condition, in Experiment 4. Bar heights reflect mean values. Points reflect individual participants. Error bars are standard error of the mean.

### Discussion

The results from Experiment 4 demonstrate that memory for word targets, unlike shapes or nonwords, benefits from a post-encoding period of wakeful rest in recall but not in recognition. This effect is similar to previous demonstrations of the wakeful rest effect (18,e.g., 21,22), but it is novel in that it extends the effect to incidentally encoded words. The current results indicate that the effect is consistent for both younger and older adults and it is observed in recall, but not recognition, performance.

## Experiment 5

In Experiment 5, we attempted to replicate the same wakeful rest effect using auditory, as opposed to visual, words as targets. We also changed the distractor task to a visual spot-the-difference task, in order to preserve minimal overlap in stimulus modality between the target and distractor materials. The purpose of this experiment was to provide an internal replication of the Experiment 4 results, as well as to test whether these effects were limited to specific stimulus domains and distractor manipulations.

### Method

#### Participants

Experiment 5 participants included a final sample of 24 younger adults and 16 older adults, recruited from the same sources described in Experiment 3. Again, on the Short Blessed Test, all older participants scored within the range of normal cognition, and so no participants were excluded for cognitive impairment.

#### Design

The design was similar to the one described in Experiment 4 with one major distinction: we tested memory for auditorily-presented words, instead of visually-presented words. Within participants, we manipulated the activity that filled the post-encoding period (either wakeful rest or a distractor task). Age group was a between-participant variable (younger or older adults).

#### Materials

Stimuli included audio recordings of the same nouns described in Experiment 4. Each recording included the word spoken aloud by a female speaker in the clear with a duration of approximately 1 second per word.

#### Procedure

The procedure was similar to the one described in Experiment 4. Aspects of the procedure were slightly changed to test the effect in auditory word stimuli, as described below.

Experiment 5 included two incidental encoding phases. During each phase, participants incidentally encoded a list of auditorily-presented nouns by providing pleasantness ratings in a task similar to the ones described in Experiments 1.

The post-encoding interval lasted 10 minutes. It consisted of either wakeful rest or a 10-minute interval of a spot the difference task, similar to the one used by Dewar and colleagues (18), etc. During the spot-the-difference task, participants viewed two pictures of the same complex scene, presented sized by side. For each picture pair, ten details had been digitally changed between the two images, including the addition or subtraction of small elements, or changes in the color, rotation, or shading of elements. Participants were given 1 minute to view each picture pair and identify those differences by mouse click. Participants viewed a total of 10 picture pairs in this manner.

After each 10-minute interval, all participants completed the same spot the difference task described above for an additional 5 minutes.

Apart from the differences described here, the procedure was identical to the one described in Experiment 4.

### Results

#### Questionnaire

Seven younger and 1 older adult reported expecting the memory test. One younger and 1 older adult reported attempting to rehearse the targets. Two participants (both younger) reported falling asleep. Each of these participants was replaced by a new participant given the same counterbalancing order of conditions.

#### Recall

Figure 10 displays the proportion of words recalled as a function of age and distractor condition. We tested the effects of healthy aging and condition on recall performance in a 2 x 2 mixed-model ANOVA, with proportion recalled as the dependent variable, age as a between-participant factor (younger or older adults) and condition as a within-participant factor (wakeful rest or spot the difference). As expected, this analysis revealed a main effect of age, *F*(1,38) = 9.07, *p* = .005, *η*_*G*_^2^ = .13, and importantly, a main effect of condition, *F*(1,38) = 6.64, *p* = .014, *η*_*G*_^2^ = .06. Again, there was no evidence of an interaction between age and condition, *F*(1,38) = 0.05, *p* = .816, *η*_*G*_^2^ < .01. As shown in Fig 10, older adults recalled fewer words than younger adults did, and both groups recalled more words in the wakeful rest condition than in the spot the difference condition.

**Fig 10.**
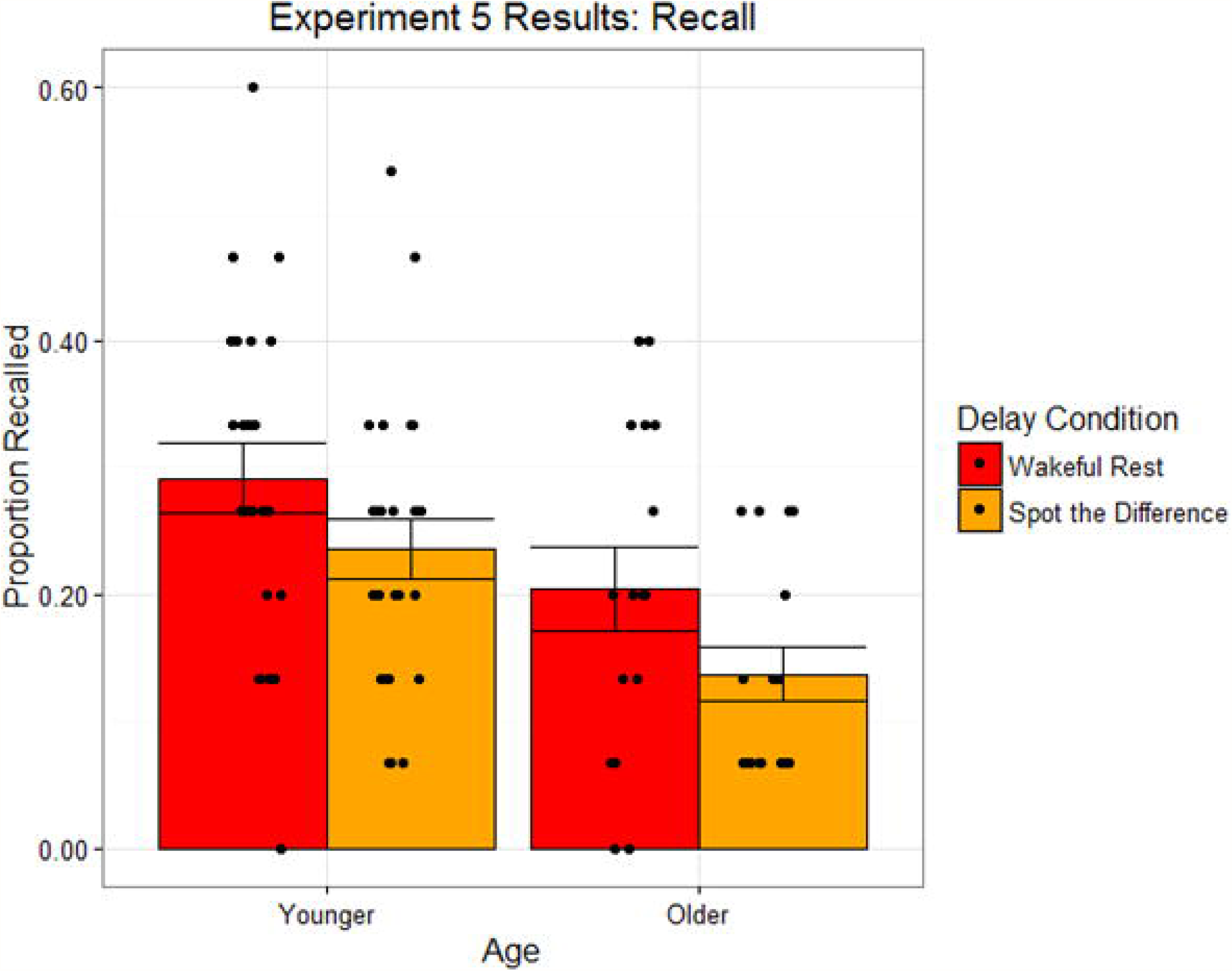
Recall task performance for auditory word targets in younger and older adults, as a function of distractor condition, in Experiment 5. Bar heights reflect mean values. Points reflect individual participants. Error bars are standard error of the mean.

#### Recognition

Figure 11 displays recognition memory performance (defined as hits-false alarms) as a function of age and distractor condition. We tested the effects of healthy aging and condition on recognition performance in a 2 x 2 mixed-model ANOVA, with the same factor structure described for recall. This analysis revealed a main effect of age, *F*(1,38) = 32.18, *p* < .001, *η*_*G*_^2^ = .41, but no main effect of condition, *F*(1,38) = 2.50, *p* = .122, *η*_*G*_^2^ = .01, or an interaction between age and condition, *F*(1,38) = 0.00, *p* = .970, *η*_*G*_^2^ < .01. Again, we followed up on the null effect of condition with a Bayesian mixed-model ANOVA. The estimated Bayes factor for the effect of condition, *BF*_*10*_, was 0.674. Hence, the data provide only anecdotal support for the null over an alternative hypothesis, *BF*_*01*_, by a factor of 1.474. As shown in Fig 11, older adults had lower recognition performance than younger adults did, and both groups performed similarly in the wakeful rest condition, as compared to the spot the difference condition.

**Fig 11.**
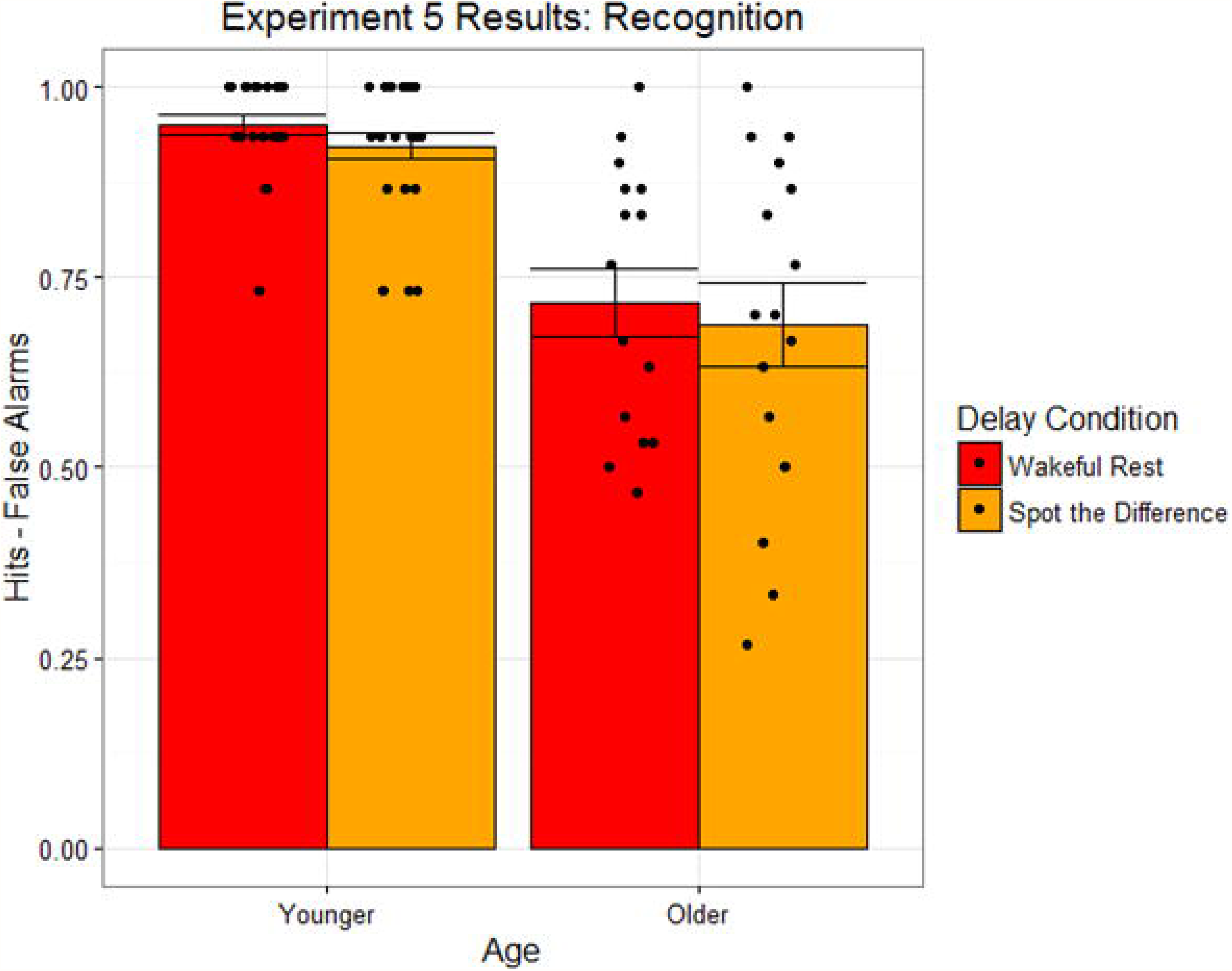
Recognition task performance for auditory word targets in younger and older adults, as a function of distractor condition, in Experiment 5. Bar heights reflect mean values. Points reflect individual participants. Error bars are standard error of the mean.

### Discussion

The results from Experiment 5 demonstrate that the wakeful rest effect for incidentally encoded words (demonstrated in Experiment 4) replicates for auditory word targets. As in Experiment 4, this effect is consistent for both younger and older adults and is observed in recall, but not recognition, performance. The failure to observe a wakeful rest effect in any recognition test (Experiments 1 – 5) is inconsistent with previous demonstrations (Craig & Dewar, 2018; Dewar et al., 2014) and poses a problem for theoretical interpretation of our results. It is possible that failures to observe a wakeful rest effect in these cases are due to a lack of sensitivity in the recognition tests we used and not some theoretical limitation on the wakeful rest effect.

## Experiment 6

In Experiment 6, we attempted to adjust the paradigm used in Experiment 4, in which we observed a wakeful rest effect in recall, but not recognition, to be more sensitive to conditional effects on recognition. Specifically, we reordered the recognition and recall tasks to place more emphasis on the recognition test, and also included a 24-hour delay between encoding and retrieval to test for conditional effects on recognition at different levels of overall performance. It is possible that the consequences of wakeful rest might develop over time, particularly during a night’s sleep. Hence, this experiment also allowed us to compare wakeful rest effects in immediate and delayed retrieval.

### Method

#### Participants

Experiment 6 participants included a final sample of 48 younger adults, recruited from the same sources described in Experiment 3. Twenty-four participants completed the memory tasks the same day as the encoding task, and 24 completed the memory tasks the following day after a 24 hour retention interval.

#### Design

The design was similar to the one described in Experiment 4 with two major distinctions: First, we reordered the memory tests in the retrieval phase such that the recognition test preceded recall. Second, in addition to the usual one-day paradigm used in Experiment 4, we also included a version with 24-hour delayed memory test. Within participants, we manipulated the activity that filled the post-encoding period (either wakeful rest or a distractor task). Testing delay was a between-participant variable (same-day or next-day).

#### Materials

Stimuli included the same nouns described in Experiment 4.

#### Procedure

The procedure was similar to the one described in Experiment 4. Aspects of the procedure were slightly changed to reorder the retrieval tests and to include a longer delayed test condition, as described below.

During the retrieval stage, all participants completed a recognition task first, which was then followed by a recall task. The tasks were identical to those described previously in Experiments 4 – 5. The only difference was the reversed ordering of the memory tasks.

Half of the participants completed the same-day testing paradigm, as described in Experiment 4. The other half completed a 24-hour delayed testing paradigm. In this version, participants ended the first day’s session after completing the final distractor task. They then returned to the lab 24 hours later to complete the surprise recognition, recall, and questionnaire tasks. In order to avoid floor performance in the 24-hour delayed test, the incidental encoding phases were also modified for these participants, in order to increase the number of encoding events. Specifically, participants in the 24-hour delayed version made two separate pleasantness ratings about each target stimulus within each encoding block.

### Results

#### Questionnaire

Thirteen participants (7 same-day and 6 delayed-test) reported expecting the memory test. Three participants (2 same-day and 1 delayed-test) reported attempting to rehearse the targets. Fourteen participants (7 same-day and 7 delayed-test) reported falling asleep. Each of these participants was replaced by a new participant given the same counterbalancing order of conditions.

#### Recognition

Figure 12 displays recognition memory performance (defined as hits-false alarms) as a function of testing delay and distractor condition. We tested the effects of delay and condition on recognition performance in a 2 x 2 mixed-model ANOVA, with the same factor structure described for recall. This analysis revealed a main effect of testing delay, *F*(1,46) = 6.83, *p* = .012, *η*_*G*_^2^ = .12, but no main effect of condition, *F*(1,46) = 0.68, *p* = .416, *η*_*G*_^2^ < .01, nor an interaction between delay and condition, *F*(1,46) = 0.03, *p* = .864, *η*_*G*_^2^ < .01. Again, we followed up on the null effect of condition with a Bayesian mixed-model ANOVA. The estimated Bayes factor for the effect of condition, *BF*_*10*_, was 0.285. Hence, the data provide substantial support for the null over an alternative hypothesis, *BF*_*01*_, by a factor of 3.509. As shown in Fig 12, the 24-hour delay group had lower recognition performance than did the same-day group, and both groups performed similarly in the wakeful rest condition, as compared to the spot the difference condition.

**Fig 12.**
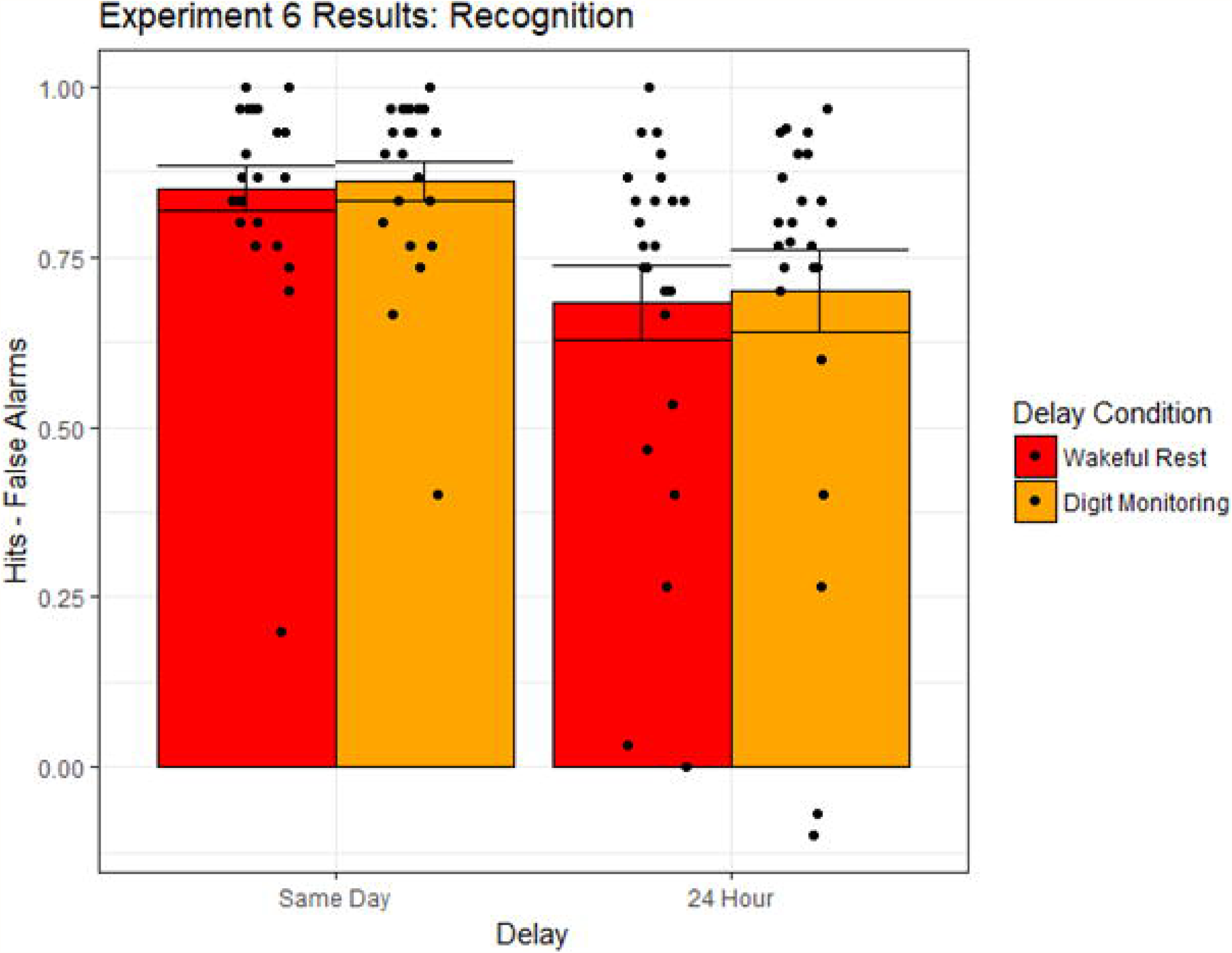
Recognition task performance for auditory word targets in younger and older adults, as a function of distractor condition, in Experiment 6. Bar heights reflect mean values. Points reflect individual participants. Error bars are standard error of the mean.

#### Recall

Figure 13 displays the proportion of words recalled as a function of testing delay and distractor condition. We tested the effects of testing delay and condition on recall performance in a 2 x 2 mixed-model ANOVA, with proportion recalled as the dependent variable, delay as a between-participant factor (same-day or delayed) and condition as a within-participant factor (wakeful rest or digit monitoring). This analysis revealed no main effect of testing delay, *F*(1,46) = 0.51, *p* = .477, *η*_*G*_^2^ = .01, no main effect of condition, *F*(1,46) = 1.70, *p* = .20, *η*_*G*_^2^ = .01, and no evidence of an interaction between delay and condition, *F*(1,46) = 0.19, *p* = .666, *η*_*G*_^2^ < .01. Again, we followed up on the null effect of condition with a Bayesian mixed-model ANOVA. The estimated Bayes factor for the effect of condition, *BF*_*10*_, was 0.479. Hence, the data provide only anecdotal support for the null over an alternative hypothesis, *BF*_*01*_, by a factor of 2.087. As shown in Fig 13, mean levels of recall did not differ significantly between the same-day group and the 24-hour delay group, nor did recall differ between the wakeful rest and distractor conditions for either group.

**Fig 13.**
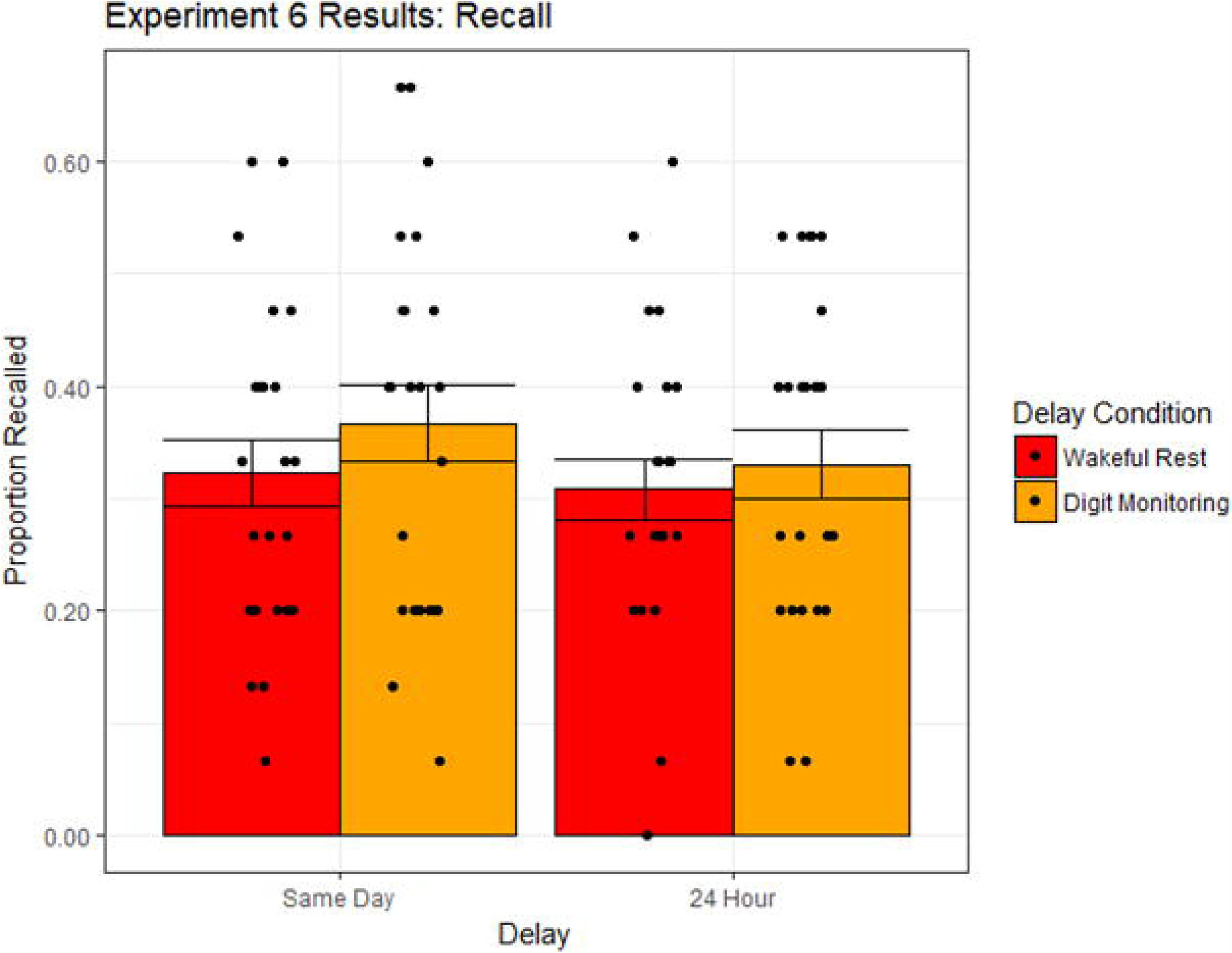
Recall task performance for auditory word targets in younger and older adults, as a function of distractor condition, in Experiment 6. Bar heights reflect mean values. Points reflect individual participants. Error bars are standard error of the mean.

### Discussion

The results from Experiment 6 once again failed to find a wakeful rest effect in a recognition test. This lack of an effect was consistent for both the same-day test and the 24-hour delayed test conditions, which produced different overall levels of performance. Hence, the overall failure to detect an effect on recognition does not appear to be due to a scaling limitation, such as a lack of sensitivity due to ceiling-level performance.

Additionally, Experiment 6 failed to replicate the wakeful rest effect in recall for incidentally encoded words (demonstrated in Experiments 4 and 5). This result, however, is not entirely surprising. Since the recall test followed recognition, participants might have been able to use the test probes from the recognition task as a chance to restudy the target words, leading to better overall performance than when recall preceded recognition. We tested for any effect of testing order by comparing recall performance in the younger adults across Experiments 4 and 6, using test order as a between-participants factor. Indeed, this analysis revealed a main effect of order, *F*(1,70) = 10.87, *p* = .002, *η*_*G*_^2^ = .13, which was further characterized by an interaction between order and condition, *F*(1,70) = 6.69, *p* = .01, *η*_*G*_^2^ = .09. Recall performance in the distractor condition was significantly greater in the recognition-first order than in the recall-first order, *t*(70) = −4.11, *p* < .001, but performance in the wakeful rest condition did not differ across testing orders, *t*(70) = −1.18, *p* = .24. Hence, the reversed testing order allowed for greater improvement in the distractor condition, eliminating the wakeful rest benefit in Experiment 6.

## General Discussion

Across all six experiments, we observed four consistent patterns. First, benefits of wakeful rest, as compared to a distractor task, were not observed for memory tests of non-verbal abstract shape targets (Experiments 1, 2, and 3). Second, significant wakeful rest effects were observed for recall of incidentally encoded words (Experiments 4 and 5, but see Experiment 6). Third, these wakeful rest effects for words were consistent for both younger and older adults (Experiments 4 and 5). Fourth, wakeful rest effects were not observed in any measure of recognition memory performance (all experiments). We now discuss each of these observations in turn, with a particular focus on how our observations fit with the previous literature and how these results inform theoretical interpretations of the wakeful rest effect.

### Failure to Extend the Wakeful Rest Effect to Abstract Shapes

Across three experiments, we found no evidence of episodic facilitation for abstract shape targets in post-encoding wakeful rest. This null effect was consistent for both incidental (Experiments 1 and 3) and intentional encoding paradigms (Experiment 2), suggesting that the intentionality of encoding does not influence the effect. Critically, these experiments differ from previous demonstrations in that the memory targets cannot be readily mediated by a verbal label. Thus, the chance of rehearsal of these memory targets during wakeful rest is much less likely, in comparison to prose passages (e.g., 18), word lists (e.g., 21), maps (e.g., 25), pictures of familiar objects (e.g., 27), or pronounceable non-words (e.g., 22), all of which can be at least verbally mediated. Hence, the current data suggest that some degree of verbal mediation may be necessary for wakeful rest to benefit recently encoded materials. This observation is predicted by a theoretical account in which the wakeful rest effect emerges as a consequence of the greater opportunities for rehearsal afforded during the wakeful rest interval. In contrast, an automatic consolidation account would make no prediction of differential effects depending on the capacity for rehearsal: both verbal and non-verbal stimuli should be equally likely to consolidate via an automatic neural replay mechanism.

Importantly, we are not the first group to report null effects in a variant of the wakeful rest paradigm. Indeed, recent failures to extend the effect have been reported for both free recall and recognition tests of verbal target materials (28,29). One account of some of these failures is provided by Varma and colleagues (28). They suggest that a distractor task should only disrupt episodic enhancement for recent encoding if the task demands episodic processing, presumably driven by the hippocampal system. They argue that the N-back working memory task, which they use as a distractor, does not engage episodic processing, and hence, does not interfere with memory consolidation in comparison to a wakeful rest period. A later experiment tested this account, directly comparing wakeful rest, a 2-back task, and an autobiographical listening task (similar to the one used presently in Experiments 1 and 2, as well as by Craig and colleagues (21)) in their effects on retention of intentionally-encoded words (40). Consistent with their earlier account, Varma and colleagues (40) only observed a disruption in retrieval performance in the autobiographical retrieval conditions, whereas wakeful rest and the 2-back task produced comparable retrieval performance. It should be noted that in this study there was also a significant relationship between self-reported stimulus-oriented thoughts, including intentional rehearsal and spontaneous thoughts about the target items, and later memory performance, which was restricted to the wakeful rest condition. However, the interpretation that the wakeful rest effect should be modulated by the episodic demands of the distractor task is not supported by the current results. In Experiments 1 and 2, we compared wakeful rest to a similar autobiographical listening task, which should presumably place a demand on autobiographical processing in the hippocampus. In Experiment 3, we used a simple 1-back digit-monitoring distractor task, similar to the n-back tasks used by Varma and colleagues (28,40), which should presumably not demand hippocampal processing. Critically, wakeful rest effects were not observed for abstract shapes (or nonwords), regardless of the distractor task selected. Hence, the failure of the wakeful rest effect in the current dataset is more likely driven by the retrieval test type and/or the target materials themselves, rather than any aspect of the distractor tasks.

In a separate account of null effects in a wakeful rest paradigm, Martini and colleagues (29) suggest that their particular encoding paradigm might have produced highly elaborated memory traces, since they gave participants unlimited time to study emotionally arousing narratives (i.e., crime and accident scenes), under unusual testing conditions (i.e., narratives were studied in participants’ second language and tested in their first language). Consequently, memory for the passages might be particularly stable and less affected by intervening distractors, limiting the sensitivity of their comparison across conditions. However, other studies have not directly tested this account. In the current study, we consistently fail to find a wakeful rest effect across a wide range of mean recognition performance levels. Hence, the elaboration or strength of a memory trace does not appear to be important to the presence or absence of the wakeful rest effect in the current study.

### Extension to Incidentally-Encoded Verbal Targets

The vast majority of previous wakeful rest experiments have used a paradigm in which verbal materials are intentionally encoded for an immediate recall test, but then memory for the same materials is also tested after a wakeful rest and/or distractor task in a surprise delayed recall test (18,e.g., 21,22). The authors have argued that, because there is an immediate recall test, participants should no longer expect a memory test, and so should be less likely to rehearse or think about the items. However, under these circumstances, the presence of the immediate recall test might signal the importance of memory for the targets, encouraging participants to engage in rehearsal of successfully-recalled targets during the subsequent wakeful rest interval. Moreover, participants might also be motivated to continue searching their memory for targets that were *not* successfully recalled in the immediate test. These behaviors could contribute to the wakeful rest memory effect through mechanisms that are rooted in executive/attentional control processes and/or controlled retrieval, rather than automatic, spontaneous consolidation. Importantly, however, the results from Experiments 4 and 5 indicate that the presence of an immediate memory test is not crucial in obtaining a wakeful rest effect in recall performance. Hence, we would argue that the presence of the immediate memory test is not necessary to finding a wakeful rest effect. In this light, it is interesting to note that Craig and Dewar (27) recently replicated the wakeful rest effect on recognition memory performance for picture targets that were encoded via a semantic incidental orienting task (i.e., indoor vs. outdoor judgments) without including an immediate memory test. Thus, it appears that the presence of an immediate memory test and the intentional encoding instructions are not critical in observing an effect in recall or recognition performance.

Regarding the distractor tasks used in these experiments, again our results are inconsistent with the suggestion by Varma and colleagues (2017, 2018). In Experiment 4, we compared wakeful rest to a 1-back digit-monitoring distractor task, which should place minimal demand on autobiographical processing. In Experiment 5, we used a spot-the-difference distractor task, similar to one used by Dewar and colleagues (18), among others, which might demand more complex processing, perhaps even involving autobiographical recall or future planning. Critically, wakeful rest effects were observed for recall of verbal targets, regardless of the distractor task selected. Hence, across all experiments using verbal and non-verbal targets, the wakeful rest effect is more likely driven by either the retrieval test type and/or the target materials themselves, rather than any aspect of the distractor tasks.

### Consistent Effects for Younger and Older Adults

In Experiments 4 and 5, not only did we find that recall of verbal targets benefited from an interval of wakeful rest, but also these benefits were consistent for both younger and older adults. This observation is consistent with previous work using a virtual map-learning task. Specifically, Craig and colleagues (25) found that younger and older adults exhibited comparable benefits of wakeful rest in map learning accuracy, as assessed using a virtual landmark pointing task. Together with the present results, these findings suggest that the processes involved in the wakeful rest memory effect might be relatively preserved in advancing age for both spatial and episodic memory systems. If this effect is indeed reflective of the consolidation stage of memory processing, this preservation would stand in contrast to many well-documented age-related deficits in other episodic memory processes (for review, see 32,33).

### Failure to Observe Effects in Recognition Memory

To our knowledge, only two previous experiments have reported significant wakeful rest effects in recognition memory performance. Specifically, Dewar and colleagues (22) found a wakeful rest effect in overall discriminability, as measured by *d’*, for both words and pronounceable nonwords. Additionally, Craig and Dewar (27) recently found that wakeful rest was also associated with increased discrimination of lures that were similar, but not identical, to incidentally encoded picture targets. In contrast to these effects, across all experiments in the current report, we found no evidence of a successful wakeful rest effect in any recognition task, regardless of the class of memory target or encoding paradigm. Instead, significant wakeful rest effects were only found in recall with verbal materials. The current task-specific benefits of wakeful rest might be expected by a theoretical account in which the wakeful rest effect is driven by strategic rehearsal, as opposed to consolidation. Specifically, verbal rehearsal of the target words during the wakeful rest interval might particularly strengthen associations across words in the target list. These intra-list associations might particularly aid performance in an unstructured free recall task, in which retrieval of one target might serve as a cue for retrieval of another target from the same list. However, in a recognition task, which tests memory for targets in a random order, intermixing multiple source contexts, these associations might minimally influence performance (see, e.g., 43).

It is also possible that methodological differences across studies might contribute to the absence or presence of a wakeful rest effect in recognition performance. Specifically, both of the previous experiments that demonstrated significant wakeful rest effects in recognition used a between-participants design (22,27). Interestingly, in each of these studies, the wakeful rest benefits on recognition performance were not driven by increased hits, but rather, by decreased false alarms to new foils (22) or increased discrimination of similar lures (27). Unfortunately, our use of a within-participants manipulation of the intervening task does not allow us to compare false alarm rates between wakeful rest and distractor conditions, as each participant only responded to one set of foil trials on the recognition task. Hence, our results suggest that wakeful rest does not produce an overall benefit in memory strength on a recognition task, but we cannot rule out the possibility that it might produce an increase in fine-grained discrimination of memory representations.

### Limitations

One general limitation of the present study is that we rely on the interpretation of null effects, particularly in the case of absent wakeful rest effects for abstract shape targets. However, each of the null effects we consider are internally replicated multiple times in independent samples across the present set of experiments. Also, as noted in the results, Bayesian analyses of the theoretically-relevant effects of condition provided substantial evidence that the observed data were more likely under the null hypothesis than under an alternative in all experiments, except Experiment 5, which only provided anecdotal evidence in favor of the null. Finally, with the exception of Experiment 1, which served as a pilot experiment, each individual sample achieved adequate statistical power to detect the wakeful rest effect. Specifically, power analyses in G*Power 3.1 (44) revealed that power to detect a medium-sized (Cohen’s *ƒ* = 0.25), within-participant main effect of distractor condition in the recognition task averaged .88 across experiments, ranging from .78 (Experiment 2) to .98 (Experiment 3). Hence, we argue that the internal replications across multiple, well-powered samples suggest that the present null effects are clear and robust.

Beyond statistical power, the failure to detect a wakeful rest effect across any of the recognition tests might also pose certain theoretical limitations. For instance, it is possible that floor or ceiling effects might obscure conditional differences in recognition performance. However, our use of multiple types of stimuli, age groups, and testing delays resulted in a wide range of mean recognition performance values. Hence, the lack of a wakeful rest effect in the present recognition tests is not likely due to scaling issues. Alternatively, it is possible that the benefits of wakeful rest might be limited to more self-generated or attentionally-controlled retrieval processes, as demanded by recall tasks. Typically, recognition is less sensitive to these processes than recall tasks (see 45), which indeed demonstrated significant wakeful rest effects in Experiments 4 and 5. Future studies might resolve this question by examining the impact of wakeful rest on process estimates of controlled and automatic retrieval processes.

### Conclusions

Previous studies using the wakeful rest paradigm have made important attempts to manipulate memory consolidation within the context of a behavioral experiment, in order to interrogate the cognitive processes involved in consolidation. However, we argue that previous work has not fully ruled out a potential rehearsal mechanism to explain the effect. In the present experiments, we made further attempts to limit the possibility of rehearsal during wakeful rest, by using incidental encoding paradigms and by testing memory for unrehreaseable abstract shapes. We found no evidence of a wakeful rest effect for these materials under a variety of distractor tasks and encoding paradigms in both young and older adults. We only found evidence for wakeful reset effects for words. Moreover, we only found such effects in a recall task that demands more explicit retrieval operations. Hence, we argue that some capacity for rehearsal is necessary to achieve the wakeful rest effect, and therefore, that these effects might not reflect a pure measure of automatic consolidation processes, independent of rehearsal.

## Acknowledgments

We thank Ben Jadow, Neco Johnson, and Jared Lassner for assistance in data collection and coding.

